# X chromosome inactivation as a novel mechanism for chondrogenesis

**DOI:** 10.64898/2026.06.23.733966

**Authors:** Hiroki Ueharu, Haichun Pan, Sher Khehra, Sundeep Kalantry, Yuji Mishina

## Abstract

X chromosome inactivation (X-inactivation) is generally regarded as a dosage-compensation mechanism restricted to female mammals. Here we show that BMP signaling induces X-inactivation through upregulation of *Xist* and promotes chondrogenesis in both sexes. Remarkably, augmented BMP signaling induced ectopic X-inactivation: transiently inactivating both X chromosomes in females and one in males in a tissue-specific manner. In cranial neural crest cells, ectopic X-inactivation suppressed X-linked gene *Tmsb4x*, leading to ectopic cartilage formation. Genetic reduction of *Xist* or pharmacological restoration of the *Tmsb4x* product suppressed this phenotype. Moreover, we identified ectopic X-inactivation in SOX9-positive chondroprogenitors during wild-type forelimb development in both sexes. Inhibition of X-inactivation disrupts proximodistal limb patterning *ex vivo*. These findings establish X-inactivation as a signaling-dependent developmental mechanism linking chromosome-scale gene alterations to skeletal fate specification.

## Main Text

Spatial and temporal regulation of cellular behaviors in multipotent cells are critical for skeletal development, including in the craniofacial region. During craniofacial development, cranial neural crest cells (NCCs), which are multipotent cells differentiating into osteoblasts, chondrocytes, and other lineages in craniofacial region, give rise to the face and the anterior part of the skull (*1–5*). Dysregulation of cranial NCCs results in craniofacial anomalies (CFAs), affecting 1 in 100 newborns (*6, 7*). Several signaling pathways have been reported to coordinate diverse cellular behaviors of cranial NCCs to orchestrate craniofacial development (*1, 8*). However, the mechanisms underlying cell fate specification remains poorly understood.

We recently reported that augmented bone morphogenetic protein (BMP) signaling in cranial NCCs in mice (*P0-Cre(+);caAcvr1-IRES-Egfp(+)*, hereafter, *caAcvr1* mut mice) develop robust ectopic cartilage in the face, particularly in the lower jaw and the tongue at mouse embryonic day 14.5 (E14.5) (*9*). Our pharmacological rescue experiments using the BMP/SMAD pathway inhibitor LDN-193189 identified stages prior to E11.5 as a critical time window for ectopic cartilage formation (*9*). These data suggest that BMP signaling regulates cell fate of cranial NCCs toward chondrogenic lineage at the early embryonic stage. The role of BMP signaling in promoting chondrogenic differentiation is well established; however, its role in regulating cell fate specification remains unclear.

X chromosome inactivation (X-inactivation) is a fundamental mechanism that equalizes X-linked gene expression between sexes of therian mammals by silencing genes on one of the two X chromosomes in females. *X-inactive specific transcript* (*Xist)* is a master regulator for X-inactivation. *Xist* is an X-linked gene that transcribes a long noncoding RNA that ‘coats’ the X chromosome from which it is expressed and recruits protein complexes, including those mediating histone methylation, to down-regulate genes on that X chromosome in a process termed X chromosome inactivation(*10–12*). Genetic upregulation of *Xist* from the second X chromosome in female cells or of the sole X in male cells can result in their ectopic inactivation (*13*); however, whether such ectopic X-inactivation occurs during embryonic organ development and what role it plays remain unknown.

In this study, we provide the first evidence that BMP signaling induces ectopic X-inactivation through upregulation of *Xist* in cranial neural crest cells (NCCs), leading to ectopic cartilage formation in *caAcvr1* mut mice of both sexes. We further demonstrate that the BMP/X-inactivation axis prompts ectopic cartilage formation through suppression of the X-linked gene *Tmsb4x*. Moreover, we identified the X-inactivation/*Tmsb4x* axis in the developing forelimb of wild-type mice in both sexes. Taken together, these findings indicate that additional X chromosome inactivation drives multipotent cells toward a chondrogenic lineage under both pathological and, unexpectedly, normal physiological conditions.

### Augmented BMP signaling in cranial neural crest cells induced ectopic *Xist* induction

Our transgenic mice expressing constitutively activated *Acvr1* in NCCs (*P0-Cre;caAcvr1-IRES-Egfp* mice, *caAcvr1* mut mice) develop ectopic cartilage in the face at E14.5 (**Fig. S1A**). Additionally, we found that *caAcvr1* mut mice develop aberrant aggregations of cells expressing SOX9, a marker gene of initiation of chondrogenic differentiation (*14*), in the first branchial arch (BA1) of embryonic day (E) 11.5 embryos (**Fig. S1C, dotted lines)** (*9*), while ectopic SOX9-aggregation was not evident in the BA1 at E10.5 **(Fig. S1B)**. Thus, we hypothesized that differentially expressed genes in cranial NCCs of *caAcvr1* mut mice at E10.5 prompts their fate to chondrogenic lineage, and then ectopic chondrogenic differentiation initiates at E11.5. To uncover the mechanisms determining the differentiation of cranial NCCs towards chondrogenic lineage, we analyzed comprehensive gene expression profiles of the BA1 cells between control mice (*P0-Cre(−);caAcvr1(+)* mice) and *caAcvr1* mut mice at E10.5 by single-cell RNA sequencing (**Fig. 1A**). The tSNE-based cell clustering of the scRNA-Seq data showed neural crest cell clusters, marked by the expression of *Hand1*/*Prrx1* (cluster 1 and cluster 2); commitment cell clusters, marked by the expression of *Hand1*/*Runx1*/*Col2* (cluster 3 and cluster 4); and a osteoblasts/chondrocytes cluster, marked by the expression of *Col1*/*Col2* (cluster 0). The ratio of osteoblasts/chondrocytes in all cells analyzed was 13.9% in control mice and 19.6% in *caAcvr1* mut mice, respectively. Conversely, the ratio of neural crest cells amongst all cells was 32.7% in control mice and 27.1% in *caAcvr1* mut mice, and the ratio of commitment cells was 21.6% in control mice and 17.9% in *caAcvr1* mut mice, respectively. These results implicated that cells of the BA1 in *caAcvr1* mut mice undergo prompted differentiation towards the osteoblast/chondrocyte lineage at E10.5. Comparison of gene expression profiles of the neural crest cell cluster (cluster 2) revealed that eleven genes exhibited significantly elevated expressions in cranial NCCs of *caAcvr1* mut mice compared to controls (**Fig. 1B**). Our previous report demonstrated that trunk NCCs in *caAcvr1* mut mice do not develop ectopic cartilage (*9*), suggesting that genes that change in expression in cranial NCCs of *caAcvr1* mut mice but not in trunk NCCs of *caAcvr1* mutant mice are candidate genes involved in ectopic cartilage formation. We excluded genes that similarly change in expression in cranial and trunk NCCs (**Fig. S2A and Fig. S2B**). This analysis yielded 11 genes that changes uniquely in cranial NCCs but not in trunk NCCs, and which we nominated as candidates for the formation of ectopic cartilage in *caAcvr1* mut mice **(Fig. 1B)**.

**Fig. 1.**
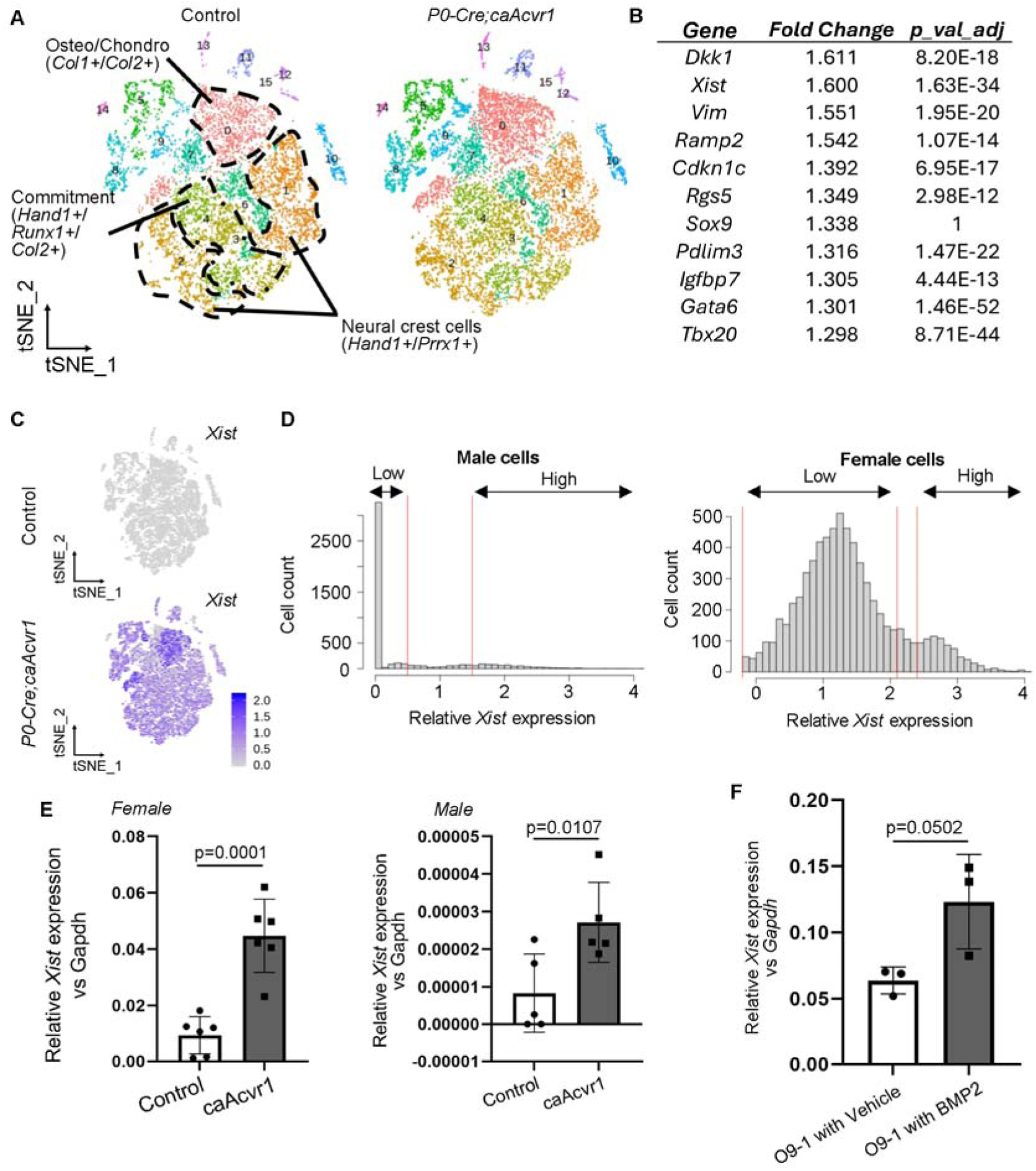
Single-cell characterization of cells of the first branchial arch at E10.5. **(A**) We prepared a single-cell suspension of the first branchial arch (BA1) of three control mouse embryos (2 males and 1 female), and similarly, from *P0-Cre;caAcvr1* mice embryos (1 male and 2 females). Unbiassed single-cell RNA sequencing for cells of the BA1 from control mice (8518 cells) and *P0-Cre;caAcvr1* mice (11834 cells) was visualized by tSNE-based analysis. The dotted area represented neural crest cell clusters (Cluster 1 and 2; *Hand 1*+/*Prrx1*+), commitment cell clusters (Cluster 3 and 4; *Hand1*+/*Runx1*+/*Col2+*), and an osteoblasts/chondrocytes cluster (Cluster 0; Col1+/Col2+). (**B**) A list of genes that expressions exhibited statistically significant changes in cluster 2 of *P0-Cre;caAcvr1* mice compared to those in control mice were shown. (**C**) Feature plots of *Xist* in control mice and *P0-Cre;caAcvr1* mice were shown. (**D**) The BA1 cells of *P0-Cre;caAcvr1* mice were further separated into male cells (3259 cells) and female cells (6944 cells). Relative *Xist* expressions in male cells and female cells of *P0-Cre;caAcvr1* mice were shown. Cells with relative *Xist* expression between 0-0.5 in males and 0.8-3.1 in females were clustered as *Xist*-low expressing cells, and cells with relative *Xist* expression over 1.5 in males and over 3.4 in females were clustered as *Xist*-high expressing cells. (**E**) Expressions of *Xist* in the first branchial arch (BA1) of control mice (white) and *P0-Cre;caAcvr1* (black) mice were quantified by quantitative reverse-transcription PCR. Female; n=6, Male; n=5. (**F**), Relative *Xist* expression in O9-1 cells with or without BMP2 (n=3 for each) was shown. The expressions were normalized with the *Gapdh* expression. Student’s *t*-test was used for statistical analysis in **E** and **F**. Each p-value is shown in the figure.

Unexpectedly, one of the eleven genes identified was the *X-inactive specific transcript* (*Xist*) (**Fig. 1B and 1C**). We next hypothesized that augmented BMP signaling ectopically induces *Xist*, followed by ectopic X-inactivation in cranial NCCs, resulting in the development of ectopic cartilage in the face. As we pooled BA1 cells from three embryos (2 males and 1 female in the control group, and 1 male and 2 females in the *caAcvr1* mut group) for single-cell RNA sequencing, we first digitally separated the BA1 cells of *caAcvr1* mut mice into female cells and male cells based on Y-linked gene expression (*Ddx3y*, *Eif2s3y*, *Kdm5d*, and *Uty*), then quantified the levels of *Xist* RNA in female and male cells, separately (**Fig. 1D and Figs. S2C-S2F**). The result suggested two categories of *Xist*-expressing cells, high and low, in both females and males (**Fig. 1D**). We confirmed elevated *Xist* RNA expression in the *caAcvr1* mut mice by RT-qPCR in female and male E10.5 embryos (**Fig. 1E**). We also confirmed that *Xist* expression is elevated by BMP-2 (100 ng/mL) treatment in O9-1 cells, a neural crest stem cell line from a female embryo (*3*) (**Fig. 1F**).

Upon induction, *Xist* RNA coats one X chromosome and silences genes in *cis* (*10–12*). We hypothesized that elevated *Xist* RNA expression induce inactivation of the second X chromosome in female BA1 cells. In cryosections of the BA1 of *P0-Cre(+);caAcvr1(+);mTmG* female embryos at E10.5 demonstrated two *Xist* RNA coats in a subset of *caAcvr1* mut nuclei (**Fig. 2A, Movie S1, and Movie S2**). Approximately 5 % of cranial NCCs in the BA1 of *caAcvr1* mut female embryos at E10.5 exhibited two *Xist* RNA coats in individual nuclei, while about 1% of cells in the BA1 of control female embryos exhibited two *Xist* RNA coats (**Fig. 2A** and **2D**). Furthermore, we found a significantly fewer cranial NCCs with no *Xist* RNA coats in a flat plane section of *caAcvr1* mut female embryos at E10.5 than those in control embryos (**Fig. 2E**). Surprisingly, we also found *Xist* RNA-coated nuclei in the BA1 of males of both control and *caAcvr1* mut mice at E10.5, and the ratio is approximately 3% in *caAcvr1* mut mice (Fig. **2B** and **2F**). About 1% of cells in the BA1 of control male embryos exhibited one *Xist* RNA coat (Fig. **2B** and **2F**). Finding of *Xist* RNA coated X chromosome in WT male BA1 cells was also unexpected and suggests that *Xist* RNA is present in male cells. Primary BA1 cells from *caAcvr1* mut mice clearly demonstrated ectopic *Xist* RNA induction in both sexes **(Fig. S3A** and **S3B)** (**Movie S3** for a female cell with 2 *Xist* RNA coats).

**Fig. 2.**
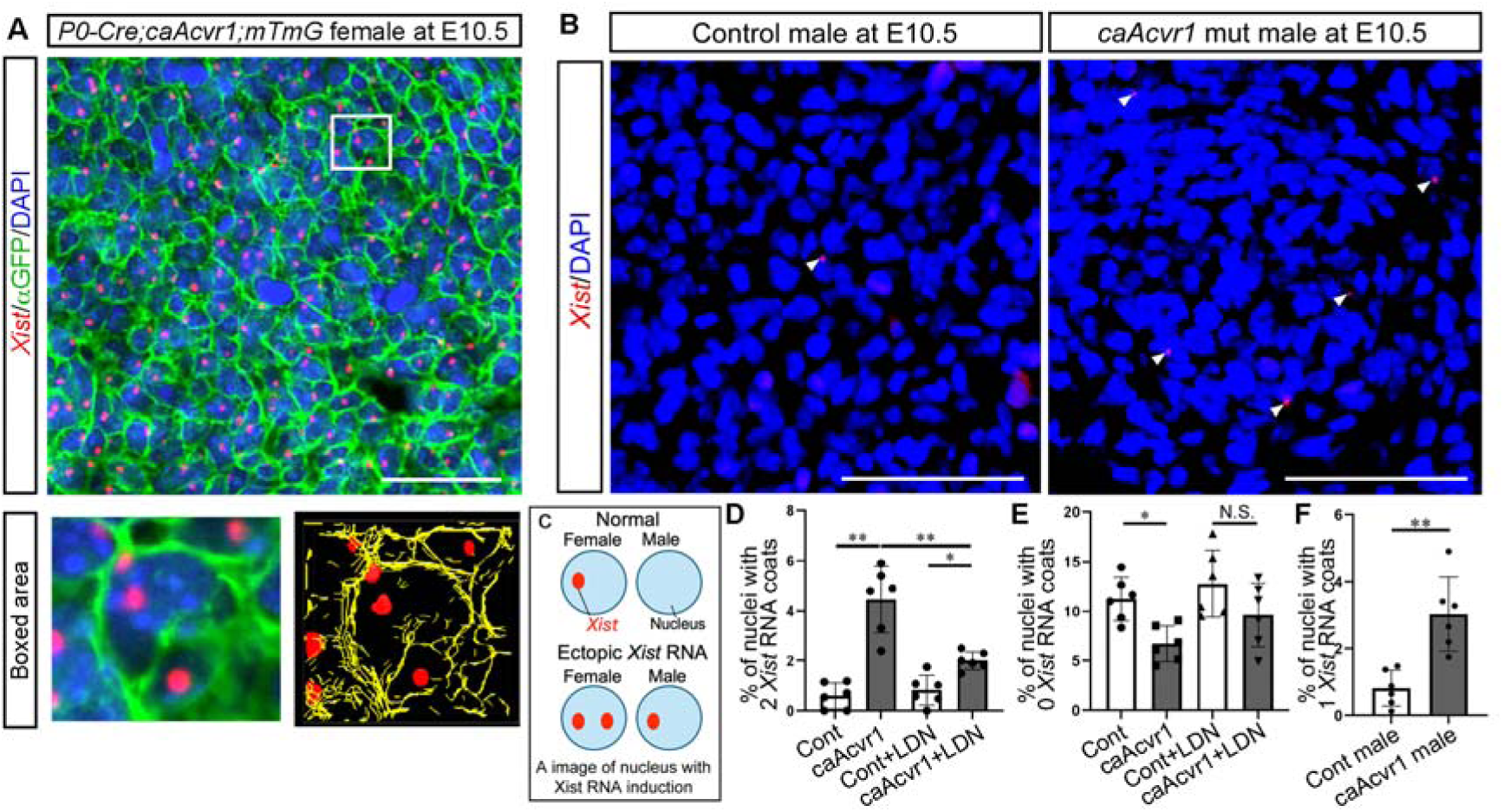
Ectopic X-chromosome inactivation induced by enhanced BMP signaling in the cranial NCCs. (**A)** Inactive X-chromosomes in the BA1 of *P0-Cre;caAcvr1;mTmG* mice were detected by *in situ* hybridization (ISH) for *Xist* RNA (Opal 570, red). Plasma cell membrane (detected by GFP antibody, Alexa 488, green) and nuclei (DAPI, blue) were visualized, respectively. The boxed area was enlarged and reconstructed to a three-dimensional image (red; *Xist*, Yellow; mGFP). **(B)** RNA scope ISH for *Xist* (Opal 570, red) in control male and *P0-Cre;caAcvr1* male at E10.5 was shown. Arrowheads indicated *Xist* RNA. **(C)** The Schematic images of a cell with ectopic X-inactivation in females and males were shown. **(D)** and **(E)**, The number of nuclei with two *Xist* RNA coats and zero *Xist* RNA coats in control female mice (cont, n=6), *P0-Cre;caAcvr1* female mice (*caAcvr1*, n=6), control female mice treated with a BMP signaling inhibitor LDN193189 (Cont+LDN, n=6), and *P0-Cre;caAcvr1* female mice with LDN193189 (*caAcvr1*+LDN, n=6) were quantified. **(F)** The number of nuclei with *Xist* RNA in control males (n=6) and *P0-Cre;caAcvr1* males (n=6) were shown. Scale bar; 50µm (**C**), 100 µm (**D**), and 10 µm (**D**, enlarged). Student’s *t*-test (**F**) and one-way ANOVA with Tukey test (**D** and **E**) were used for statistical analyses. *=p<0.05, **=p<0.01, N.S.=no significance.

We next sought to test whether downregulating BMP signaling in cranial NCCs would rescue the ectopic *Xist* RNA expression in *caAcvr1* mut mice. Pharmacologic inhibition of BMP/SMAD signaling using a selective BMP signaling inhibitor LDN193189 (LDN) (*15*) resulted in a lower number of cranial NCCs with two *Xist* RNA coats in the BA1 of *caAcvr1* mut female embryos compared to those in the BA1 of *caAcvr1* mut female embryos without LDN treatment (**Fig. 2D**). The number of 0 *Xist* RNA coats in *caAcvr1* mut mice showed similar level to the control upon treatment with the BMP signaling inhibitor LDN193189 (**Fig. 2E**). Together, these data indicate that BMP signaling is both necessary and sufficient to induce *Xist* RNA.

### BMP-induced ectopic *Xist* induction contributes to ectopic cartilage formation

We next investigated whether ectopic *Xist* induction in cranial NCCs contributes to ectopic cartilage formation. The X-linked gene *Kdm5c* positively regulates *Xist* expression through its histone demethylation activity (*12*). To suppress ectopic *Xist* RNA induction in the BA1 cells of *caAcvr1* mut mice, we superimposed a *Kdm5c* flox allele in *caAcvr1* mut mice (*P0-Cre;caAcvr1;Kdm5c^fl/+^* mice and *P0-Cre;caAcvr1; Kdm5c^fl/Y^* mice). The expression of *Xist* in the BA1 was significantly lower in *P0-Cre; caAcvr1;Kdm5c^fl/+^* female mice compared to *caAcvr1* mut female mice (**Fig. S4A**). The expression of *Xist* in the BA1 of male mice also showed the similar pattern between the three genotypes (**Fig. S4A**). The number of cranial NCCs with two *Xist* RNA coats in sections was significantly lower in *Kdm5c* cKO embryos at E10.5 compared to *caAcvr1* mut female embryos at E10.5 (**Fig. 3A, and 3B**). Conversely, the number of nuclei with no *Xist* RNA coats in sections was significantly higher in *Kdm5c* cKO female mice compared to the control and *caAcvr1* mut mice (**Fig. 3B**), suggesting that deletion of one copy of *Kdm5c* in NCCs partially suppresses induction of *Xist* RNA. The level of phosphorylated-SMAD1/5/9 (p-SMAD1/5/9) was comparable between *caAcvr1* mut mice and *Kdm5c* cKO female mice, whereas there were few cells with p-SMAD1/5/9 in control mice (**Fig. 3C and Fig. S4B**). Hence, regulation of *Xist* expression by KDM5C was independent of BMP/SMAD pathways. As we previously reported (*9*), all *caAcvr1* mut mice developed midfacial defects (**Fig. 3E and 3F**), including deformation of the Meckel’s cartilage (**Fig. 3F, arrows**) along with the ectopic cartilage in the face (arrowheads in **Fig. 3F** and in **Fig. S4C**). *Kdm5c* deletion partially restored the craniofacial region along with a reduction of ectopic cartilage formation (**Fig. 3E and 3F**), remarkably, in both males and females. KDM5C regulates many genes through histone demethylation (*16*). One potential concern was that changes in the altered gene expressions by deletion of *Kdm5c* may result in inhibiting ectopic cartilage formation in *caAcvr1* mut mice. However, craniofacial development, including nasal cartilage development, between *P0-Cre(+); Kdm5c^+/+^* mice and *P0-Cre(+); Kdm5c^fl/+ or^ Kdm5c^fl/Y^* mice appeared normal (**Fig. S4D**). These results suggested that suppression of ectopic *Xist* RNA induction in the face of *Kdm5c* deficient *caAcvr1* mut mice is the primary reason for the absence of ectopic cartilage, rather than the other functions of KDM5C during the cartilage development in the face. These data support the hypothesis that induction of ectopic X-inactivation by elevated *Xist* expression leads to chondrogenic differentiation.

**Fig. 3.**
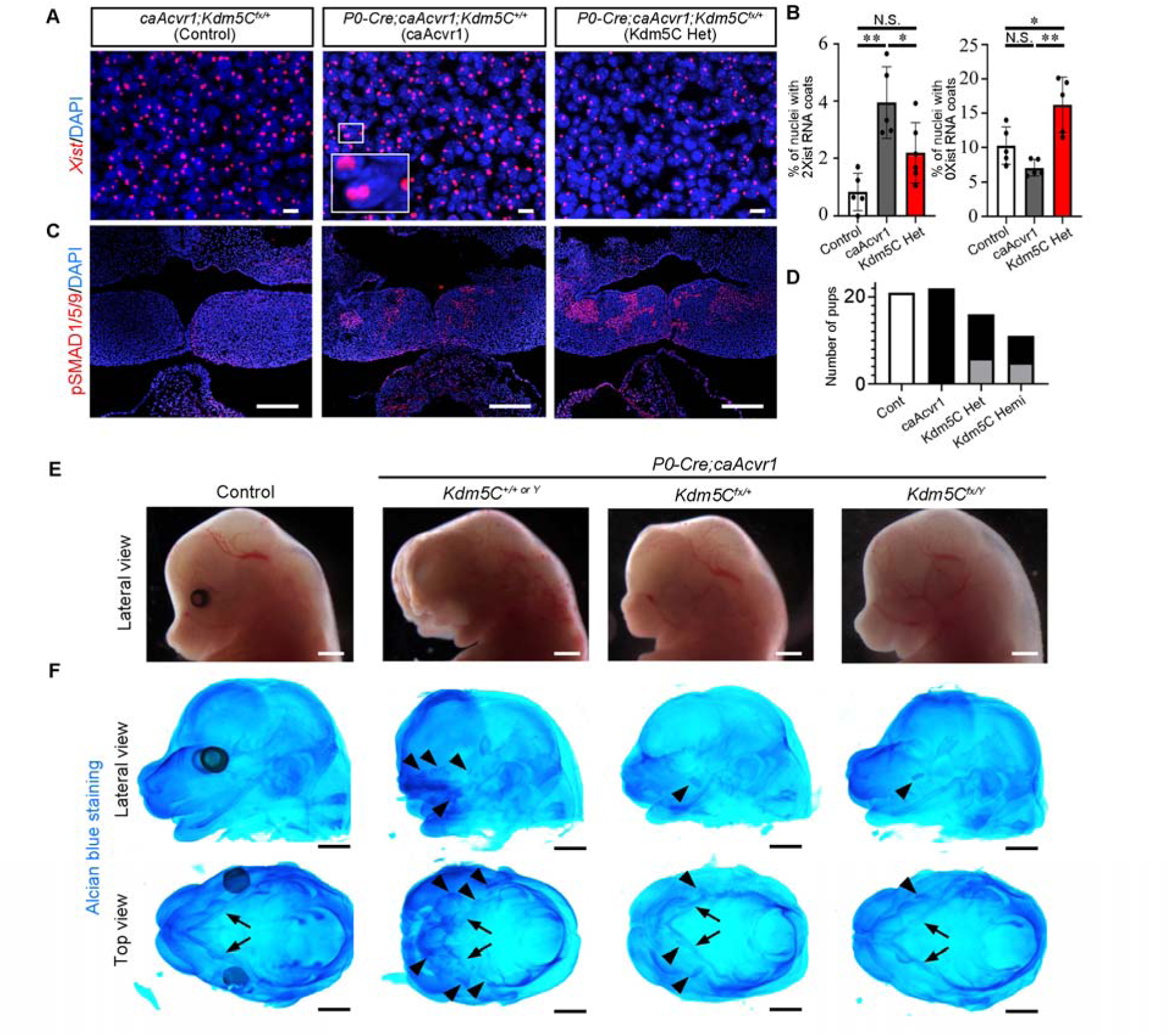
Genetic reduction of *Xist* expression in *P0-Cre;caAcvr1* mice partially rescued the midfacial defects along with reduction of ectopic cartilage. **(A)** RNA scope *in situ* hybridization for *Xist* (Opal 570, red) in the BA1 at E10.5 of *caAcvr1;Kdm5c^fl/+^*mice (control), *P0-Cre;caAcvr1;Kdm5c^+/+^* mice (*caAcvr1*), and *P0-Cre;caAcvr1;Kdm5c^fl/+^* mice (*Kdm5c* Het cKO) were shown. Nuclei were stained with DAPI (blue). The boxed area showed a nucleus with two *Xist* RNA coats. **(B)** The numbers of nuclei with two *Xist* RNA coats and nuclei with zero *Xist* RNA coats in each genotype were manually counted (n=5). **(C)** Cells with activated BMP signaling were revealed by immunohistochemistry for phosphorylated-SMAD1/5/9 (Alexa 594, red). **(D-E)**, Representative images for each genotype at E14.5 were shown in (**E**). Whole cartilage staining revealed the Meckel’s cartilage (arrows) and ectopic cartilage (arrowheads) in the face at E14.5 (**F**). The numbers of each phenotype (normal; white, severe; black, and mild; gray) at E14.5 were shown in (**D**). Control (n=20), *caAcvr1* (n=22), *Kdm5c* Het female (n=16), and *Kdm5c* Hemi male (n=11). One-way ANOVA with Tukey test (**B**) were used for statistical analyses. Scale bar=10 µm (**A**), 100 µm (**C**), and 1 mm (**E** and **F**). *=p<0.05, **=p<0.01, N.S.=no significance.

### Suppression of X-linked gene *Tmsb4x* results in ectopic cartilage formation

Next, we aimed to identify X-linked genes involved in ectopic cartilage formation. To identify X-linked genes of which expression is suppressed by ectopic *Xist* induction, we compared gene expression profiles of the *Xist*-high expressing cells and the *Xist*-low expressing cells (**Fig. 4A** and **4B**), obtained from scRNA sequencing in **Fig. 1D**. We identified 57 X-linked genes with a statistically significant reduction in the *Xist*-high expressing cells compared to the *Xist*-low expressing cells (**Fig. 4B**). Among them, we focused on the X-linked gene *Tmsb4x* (encodes Thymosin beta 4; Tβ4), which showed the most robust reduction in the *Xist*-high expressing cells. Detection of Tβ4 by immunohistochemistry (IHC) revealed that Tβ4-positive cells were broadly localized in the tongue of control mice (**Fig. 4C**). As expected, Tβ4-positive cells in *caAcvr1* mut mice was reduced in EGFP-expressing (= high BMP signaling) NCCs, and the expression of Tβ4 in EGFP-expressing cells in *Kdm5c* deficient *caAcvr1* mut mice was restored (**Fig. 4C**). We also tested Tβ4 expression during chondrogenic differentiation in NCCs using neural crest stem cell line O9-1 cells (*3*). When O9-1 cells were cultured in a pellet condition with basal media containing FGF2 and LIF to maintain their stemness, about 35 % of cells in the pellet were positive for Tβ4 (**Fig. S5A**). In contrast, a significant reduction of Tβ4-positive cells was observed when pellets were cultured in chondrogenic differentiation media (**Fig. S5A**). Moreover, culturing O9-1 cells by chondrogenic differentiation media with Tβ4 reduced the number of COL2-positive cells *in vitro* (**Fig. S5B**). These findings suggest that the reduction of *Tmsb4x* expression is associated with chondrogenic differentiation. We further tested the impact of Tβ4 by administration of Tβ4 in gestating embryos, which resulted in partially rescued midfacial region along with the absence of ectopic cartilage formation (47% of females and 56% of males were partially rescued), while all vehicle-treated *caAcvr1* mut mice showed the midfacial defects with ectopic cartilage in the face (**Fig. 4C**). These data demonstrated that BMP-induced ectopic *Xist* RNA induction develops ectopic cartilage through the reduction of the X-linked gene *Tmsb4x*.

**Fig. 4.**
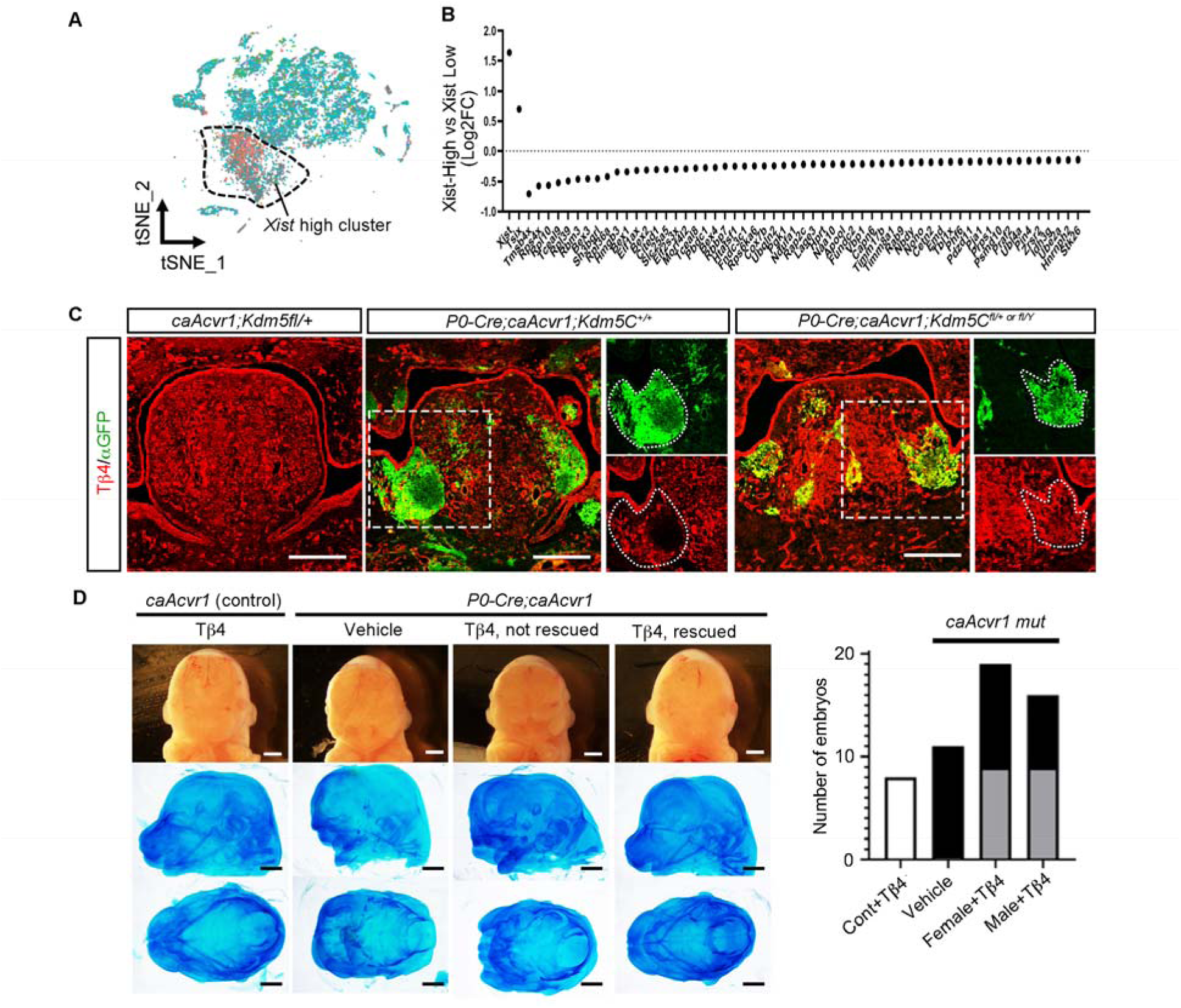
Reduction of X-linked gene Tmsb4x develops ectopic cartilage. **(A)** The comprehensive gene expression profiles of BA1 cells from *P0-Cre;caAcvr1* mice at E10.5, obtained through single-cell RNA sequencing, were re-clustered to *Xist*-high expressing cells in female (orange), *Xist*-low expressing cells in female (blue), *Xist*-high expressing cells in male (green), *Xist*-low expressing cells in male (purple), and others (gray). **(B)** X-linked genes that showed statistically significant changes in the *Xist*-high cluster compared to the *Xist*-low expressing cells in caAcvr1 mut females were shown. **(C)** Expression of Tmsb4x (Tβ4, red) in *caAcvr1;Kdm5c^fl/+^* mice (control), *P0-Cre;caAcvr1;Kdm5c^+/+^*mice, and *P0-Cre;caAcvr1;Kdm5c^fl/+^* mice was revealed by immunohistochemistry (IHC) for Tβ4. Neural crest cells expressing *caAcvr1* were visualized with IHC for GFP (αGFP, green). Nuclei were visualized with DAPI (blue). The boxed areas were enlarged on the right side. **(D)** Representative images for Tβ4 treated embryos at E14.5 were shown. The pregnant mothers intraperitoneally received Tβ4 peptide from E10.5 to E13.5. The number of each phenotype (normal; white, severe; black, and mild; gray) were shown. Control+Tβ4 (n=8), *caAcvr1* mut with vehicle (n=11), *caAcvr1* mut female with Tβ4 (n=19), and *caAcvr1* mut male with Tβ4 (n=16). Scale; 100 µm (**b**) and 1 mm (**c**).

### Ectopic *Xist* induction in the developing limbs in wild-type mice

Our next focus was to reveal the biological significance of ectopic *Xist* induction. Here, we aimed to identify the tissues that are capable to induce *Xist* ectopically by BMP signaling during normal development. Because trunk NCCs of *caAcvr1* mut mice do not develop ectopic cartilage (*9*), we hypothesize that trunk NCCs are not capable of inducing *Xist* ectopically. As we expected, we did not observe any statistical changes in the number of cells with ectopic *Xist* induction in the dorsal root ganglion between control and *caAcvr1* mut mice (**Fig. S6A**). In addition, there were no statistical differences in *Xist* expression between trunk NCCs of control and *caAcvr1* mut mice, as revealed by scRNA sequencing (**Fig. S2A**). These data suggested that BMP-induced ectopic *Xist* induction occurs in a tissue-specific manner. To identify other tissues that have the potential to induce *Xist* ectopically by BMP signaling, we next enhanced BMP signaling in whole embryos by using *Ubiquitin-CreER (+);caAcvr1(+)* mice (hereafter, *Ubc-CreER;caAcvr1* mice). First, we induced Cre-mediated recombination by tamoxifen administration at E8.5 (condition A), and then harvested the BA1, forebrain, nasal process, heart, forelimb, and trunk at E11.5. As expected, the BA1 showed a significant upregulation of *Xist*, and we also found an elevated *Xist* expression in the forelimb at E11.5 (**Fig. S6B**). Other tissues, the forebrain, the nasal process, the heart, and the trunk tend to have a tendency of a higher *Xist* expression in *Ubc;caAcvr1* mice compared to control mice, but these changes were not statistically significant. We further found that male cells of the BA1 and the fore limb in condition A showed a higher number of cells with *Xist* RNA coats in *Ubc;caAcvr1* mice compared to control mice, but not in the nasal process and the Trunk (**Fig. S7**). On the other hand, *Xist* expression at E12.5 in *Ubc;caAcvr1* mice with tamoxifen-inducible Cre recombination at E9.5 (condition B) did not show any significant differences in all tissues, including the BA1 and the forelimb (**Fig. S6C**). These data suggested that BMP signaling upregulates *Xist* expression in a spatial and temporal-specific manner.

Through this analysis, we noticed that control male cells in the forelimb at E11.5 exhibited a robust number of nuclei with *Xist* induction (**Figs. S7C and S7E**). Based on these data, we sought a possibility that ectopic X-inactivation may occur under normal physiological conditions in the forelimb of wild-type (WT) mice at E11.5. Control mice at E11.5 showed the condensed SOX9-expressing mesenchymal cells, which is the initiation of chondrogenic differentiation in the forelimb (**Fig. 5A**, boxed area) and the condensation sites for the rib cage (**Fig. 5A**, arrowheads). The presence of Tβ4 was significantly downregulated in SOX9-expressing cells (**Fig. 5B**), which is a consistent observation found in *P0-Cre(+);caAcvr1(+)* embryos (**Fig. 4C**). RNA scope *in situ* hybridization for *Xist* revealed that 12.2 % of SOX9-expressing cells showed ectopic induction of *Xist* RNA-coated X chromosomes in the forelimb of WT female at E11.5, which is a higher ratio compared to 4.2 % in SOX9-negative cells (**Fig. 5C**). We further revealed that 34.4 % of SOX9-expressing cells showed *Xist* RNA-coated X chromosomes in WT male forelimbs at E11.5 compared to 6.7 % in SOX9-negative cells (**Fig. 5D**). These data suggest that cells in the forelimb of both sexes additionally induce *Xist* RNA during normal embryonic development.

**Fig. 5.**
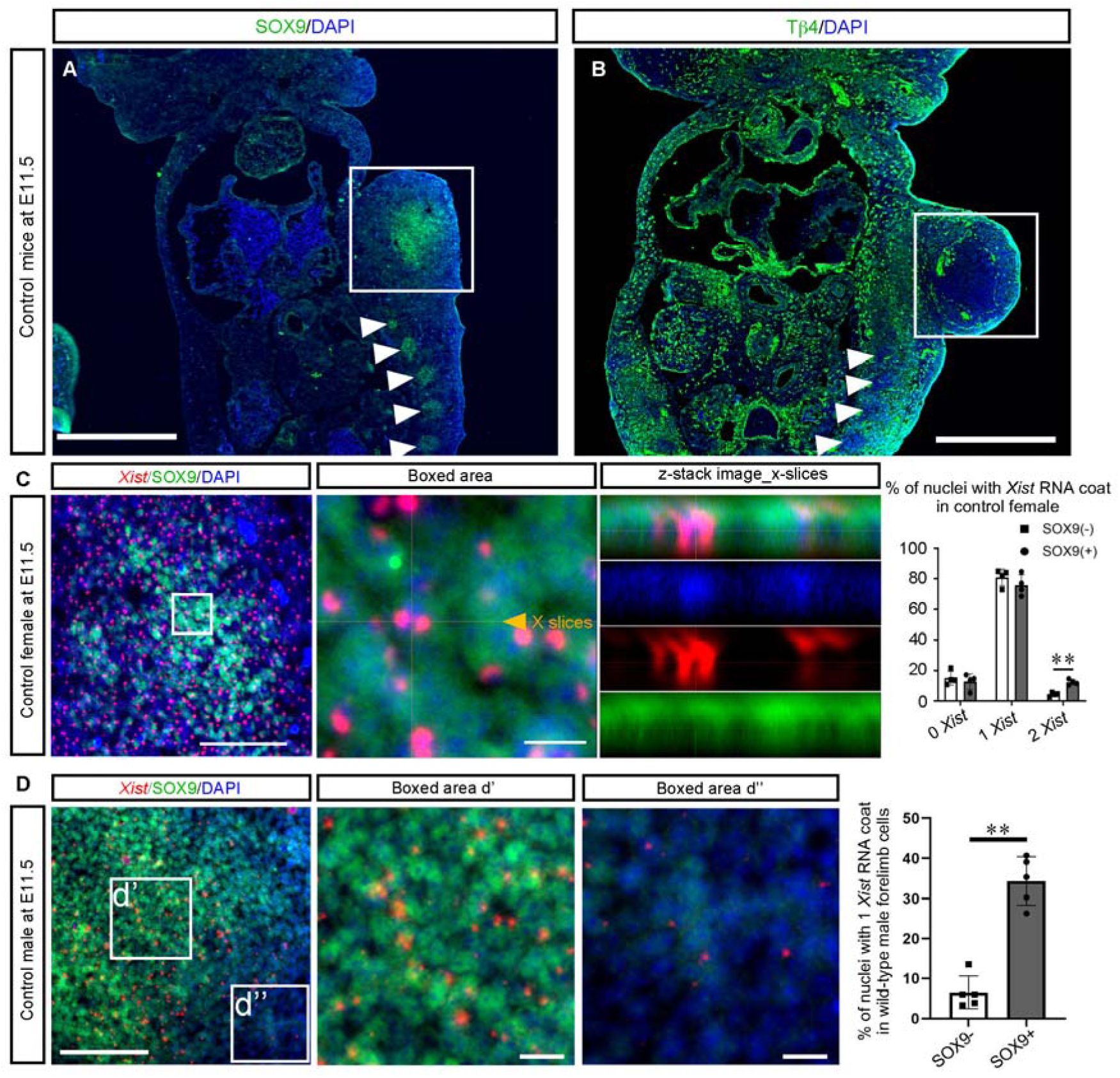
Ectopic X-inactivation in the developing forelimb. **(A-B)**, Localization of SOX9-expressing cells (**A**, n=4) and Tβ4-expressing cells (**B**, n=4) in control male mice at E11.5 are shown. White boxes showed the forelimb. Arrowheads indicated the SOX9-expressing cell clusters at the trunk region. **(C)** Inactive X-chromosomes (*Xist*, red) and chondroprogenitor cells (SOX9, green) in the fore limb of control female mice at E11.5 were shown. The boxed area was enlarged and generated the z-stack image to show 2 *Xist* RNA-coated nucleus. The number of nuclei with 0 *Xist* RNA coats, 1 *Xist* RNA coats, and 2 *Xist* RNA coats in SOX9-negative cells (white bar, SOX9 (−) cells) and SOX9-expressing cells (gray bar, SOX9(+) cells) were manually counted (n=4). **(D)** Inactive X-chromosomes (*Xist*, red) and chondroprogenitor cells (SOX9, green) in the fore limb of control male mice at E11.5 were shown. The boxed area in **(D)** was enlarged in (**d’)** and (**d’’)**, respectively. The number of nuclei with 1 *Xist* RNA coats in SOX9(−) cells and SOX9(+) cells were manually counted. Scale bars; 500 µm (**A** and **D**), 100 µm (**C and D**), and 10 µm (enlarged in **C,** enlarged in **D**). Student’s *t*-test was used for statistical analyses. *=p<0.05, **=p<0.01.

To reveal the impact of the additional *Xist* induction on forelimb development, we evaluated the patterning of forelimb by *ex vivo* organ culture. First, we cultured embryonic forelimbs *ex vivo* with X1 compound. X1 compound inhibits X-inactivation by binding to *Xist* RNA and preventing the recruitment of histone modifiers (*17*), e.g., Polycomb repressive complex 2 (PRC2)-catalyzed trimethylation of lysine 27 of histone H3 (H3K27me3). An *ex vivo* culture for WT female forelimbs with vehicle for 5 days showed that most of *Xist* RNA coats overlapped with H3K27me3 enrichment on histological specimen (16 μm thickness) (**Fig. S8A**). About 4.4 % of nuclei in female forelimbs has 2 *Xist* RNA coats overlapped with H3K27me3 enrichment, and 0.3 % of nuclei has 2 *Xist* RNA coats, one with and one without H3K27me3 enrichment (**Fig. S8B**). *Ex vivo* culture of WT female forelimbs with X1 compound, in contrast, reduced H3K27 methylation: the ratio of 2 *Xist* RNA coats overlapped with H3K27me3 enrichment was reduced to 1.5 %, the ratio of 2 *Xist* RNA coats, one with and one without H3K27me3, was elevated to 2.2 %. Moreover, nuclei with 1 *Xist* RNA coats without H3K27me3 (0.3%) and nuclei with 2 *Xist* RNA coats without H3K27me3 (0.3 %) appeared when we cultured them with X1 compound. On the other hand, an *ex vivo* culture for WT male forelimbs with vehicle for 5 days exhibited 2.8 % of nuclei with 1 *Xist* RNA coats with H3K27me3 enrichment (White arrowheads) and 8.0 % of nuclei with 1 *Xist* RNA coats without H3K27me3 enrichment (orange arrows) (**Fig. S8C**). When male forelimbs from WT mice were cultured with X1 compound, the ratio of nuclei with 1 *Xist* RNA coats with H3K27me3 enrichment was reduced to 0.8 %, and the ratio of nuclei with 1 *Xist* RNA coat without coincident H3K27me3 enrichment was elevated to 10.9 %, respectively. As expected, the X1 compound suppressed H3K27me3 on X chromosomes associated with *Xist* RNA induction during forelimb *ex vivo* culture in both sexes, while minimal impact on the *Xist* coating process on X chromosomes.

Next, we performed fore limb *ex vivo* culture by utilizing *HoxA11-GFP* mice (*18, 19*) to visualize the proximo-distal patterning of the forelimb. During normal embryonic development, *HoxA11-GFP* expressing cells are broadly expressed in the forelimb at E10.5, subsequently, limited to the developing zeugopod region (**Fig. 6A**) (*18, 20*). An *ex vivo* culture of the forelimb started at E10.5 with vehicle for 5 days showed that *HoxA11-GFP* cells localized to the middle region of the tissue, displaying a pattern similar to that of normal development (**Fig. 6B**). However, an *ex vivo* culture of the forelimb at E10.5 with X1 compound for 5 days showed the higher GFP intensity in both the middle region (ROI #1) and the proximal region (ROI #2) (**Fig. 6B and 6C**). Histological sections of forelimbs cultured with X1 compound further revealed an elevation of the number of GFP-positive cells in mesenchymal-like tissue in the proximal region compared to controls (**Fig. 6D**). Notably, GFP-positive cells in the proximal region were positive for the X-linked Tmsb4x (Tβ4), and the ratio of Tβ4+GFP+ in total area was elevated when the forelimbs were cultured with X1 compound (**Fig. 6D**). These data suggest that additional *Xist* RNA induction in normal forelimb development regulate the forelimb patterning through suppression of X-linked genes in a spatial and temporal manner.

**Fig. 6.**
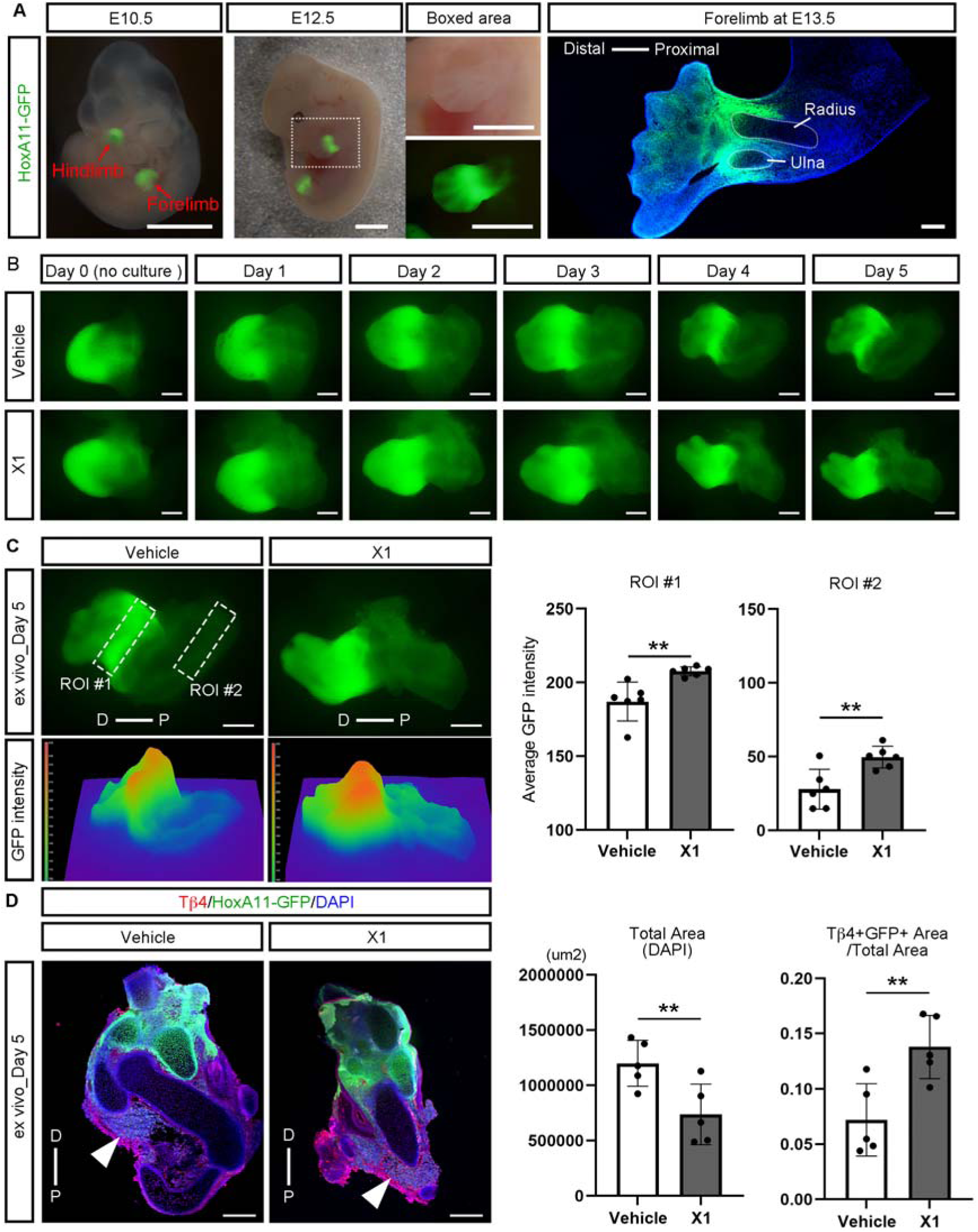
The impact of X-inactivation for the forelimb patterning. **(A)** Localization of *HoxA11-GFP* cells at E10.5 (n=4), E12.5 (n=4), and E13.5 (N=4) were shown. **(B)** Representative images of the *ex vivo* culture for the forelimb started at E10.5 with vehicle or 10 µM X1 compound for 5 days were shown. The fluorescence of *HoxA11-GFP* was observed under live imaging, every day. **(C)** GFP intensity in the forelimbs cultured for 5 days were quantified, and average GFP intensity was shown. The region with high GFP fluorescence intensity, presumed to correspond to the zeugopod, was defined as region of interest (ROI) #1, and the proximal region with low GFP fluorescence intensity was defined as ROI #2 (n=6). D; distal, and P; proximal. **(D)** Localizations of Tβ4 and HoxA11-GFP was revealed using histological sections of cultured forelimbs. Tβ4 and HoxA11-GFP double positive cells were indicated by arrowheads. D; distal, and P; proximal. Scale bars; 1 mm (**A**), 200 µm (**B**, **C**, **D**).

These data suggest an unappreciated mechanism that a spatial and a temporal expression of *Xist* in both male and female forelimbs contributes to limb patterning through suppression of X-linked genes.

## Discussion

Here, we propose a novel mechanism for chondrogenesis, that is, the BMP-induced ectopic X-inactivation. X-inactivation is believed to be a cell-autonomous mechanism occurred only in placental mammal females, and thus it had never been thought to be involved in morphogenesis. On the other hand, BMP signaling, as an extracellular signaling, largely involves in morphogenesis of both sexes, and thus the relation between BMP signaling and X-inactivation has not been focused. Only a few reports showed that the BMP/SMAD pathway directly regulates *Xist* expression (*21, 22*); however, the roles of the BMP/X-inactivation during morphogenesis had been never explored. Here, we provide the first evidence of X-inactivation contributing to chondrogenesis and normal skeletal development: A tissue-specific and transient X-inactivation is an unappreciated mechanism for skeletogenesis in both sexes.

Ectopic X-inactivation, as an additional inactivation due to elevated *Xist* expression, has been observed in mice lacking the *Xist*-antisense repressor *Tsix* (*13*). However, cells with ectopic X-inactivation disappeared from embryos of *Tsix* KO mice by E9.5 (*13*), likely due to suppression of X-linked gene expressions that are essential for cell survival. Interestingly, our *caAcvr1* mutant mice exhibited cells with ectopic X-inactivation (**Fig. 2**), but *caAcvr1* mutant mice did not elevate the number of dying cells compared to controls (*9*), likely due to the transient and tissue specific augmentation of *Xist* expression (**Figs. S6, S7**). A comprehensive analysis of inactivated X-linked genes between 27 different tissue types revealed X-linked genes subjected to X-inactivation and X-linked genes that can escape from X-inactivation (*23*). Among over 1,000 genes on X-chromosomes, we found 57 X-linked genes of which expressions were statistically downregulated in *Xist*-high expressing cells compared to *Xist*-low expressing cells in *caAcvr1* mut mice. Interestingly, within the 57 X-linked genes, 8 of them, including *Tmsb4x*, were reported as X-inactivation escapee genes (*23, 24*). These data suggested that a mechanism of ectopic X-inactivation in cranial NCCs of *caAcvr1* mut mice and SOX9-expressing chondroprogenitors in the developing forelimb may be different from the regular random X-inactivation for dosage compensation, and thus these cells can survive to demonstrate alternative behaviors. Clarifying chromatin accessibilities between random X-inactivation and ectopic X-inactivation will further reveal the function of ectopic X-inactivation during skeletal development.

This study provides the initial evidence that X-inactivation contributes to skeletal development, and thus provides a possibility in mis-regulation during the transient and tissue-specific ectopic X-inactivation may lead to skeletal disorders in humans. In general, X-linked diseases show gender biases and usually show severe phenotypes in males because females could inactivate mutant alleles in half of total cells by the regular random X-inactivation (*25*). Importantly, our *caAcvr1* mut mice induced ectopic X-inactivation in both females and males and there are no obvious phenotype differences between sexes. We further reveal that SOX9-positive cells at the forelimb bud at E11.5 induce additional X-inactivation in WT males and females. SOX9 is largely known as one of the major downstream targets of BMP signaling during chondrogenesis in both sexes (*26*). Notably, SOX9-positive cells are negative for Tβ4 (**Fig. 5B**). These data suggest that mesenchymal cells receiving BMP signaling upregulate *Sox9*, and in parallel, suppress *Tmsb4x* through additional X-inactivation, thereby initiating chondrogenic differentiation under normal physiological condition in both sexes. Therefore, we propose that additional X-inactivation plays a critical role in chondrogenesis in both sexes, and abnormal BMP/X-inactivation can cause skeletal deformities in both sexes similarly.

It is an interesting question why placental mammals downregulate X-linked genes during cartilage development through additional X-inactivation. One possible explanation is that chondrogenic gene clusters are evolutionally translocated to X chromosomes, and mammals repurposed the X-inactivation mechanism, which was originally developed for dosage compensation, to suppress those gene expressions to ensure proper skeletogenesis. Indeed, chromosome X in humans correlates with chromosome 4, 7, and X in opossums (*27*), belonging to marsupials who have evolved along with placental mammals. Interestingly, chromosome 4 and 7 in opossum code orthologs of *Tmsb4x*, *Ofd1* (a causative gene for Orofaciodigital syndrome type 1 (*28*)), *Utx*/*Kdm6a* (a causative gene for Kabuki syndrome (*29*)), and *Amelx* (a causative gene for X-linked Amelogenesis Imperfecta (*30*)), suggesting that gene clusters critical for skeletogenesis evolutionally translocated to X-chromosomes in human. Comparative gene expression profiles of X-linked genes and their ortholog during skeletal development between several species would provide new insights regarding the BMP/X-inactivation/X-linked genes during morphogenesis.

## Materials and Methods

### Mouse breeding

Transgenic mouse line carrying the Cre-inducible constitutively activated *Acvr1* (*caAcvr1*) was described previously (*31*). We then crossed the *caAcvr1* mice with *P0-Cre* mice (C57BL/6JTg(P0-Cre)94Imeg (ID 148) provided by CARD, Kumamoto University, Japan) (*32*) to generate mice with neural crest specific enhanced BMP signaling. To distinguish single cells on histological sections, we further superimposed *R26^mTmG^* (B6.Cg-Gt(ROSA)26Sortm14(CAG-tdTomato)Hze/J, The Jaxon Laboratory, #007914) in *P0-Cre;caAcvr1* mice. *Kdm5c* flox mice (*Kdm5c^tm1.2Yshi^*) (*16*) were also superimposed with *P0-Cre;caAcvr1* mice to perform the genetic rescue experiments. *HoxA11-GFP* mice (*Hoxa11^tm1Dmwe/J^*, the Jaxon Laboratory #011036) were directly obtained from Dr. D. Wellik (*18*). The day of when vaginal plugs were found was designed as embryonic day 0.5 (E0.5). All mice were housed in cages in a 20 °C room with a 12-hours (h) light/dark cycle. All animal experiments were performed in accordance with the policy and federal law of judicious use of vertebrate animals as approved by the Institutional Animal Care and Use Committee (IACUC) at the University of Michigan (Protocol #PRO00011263).

### Pharmacologic treatments for pregnant mice

The BMP receptor kinase inhibitors LDN-193189 were kindly provided by RIKEN. A dose of 2.5mg of the LDN-193189 per kg body weight was injected intraperitoneally into pregnant mice on E9.5, then embryos were harvested at E10.5 (*9, 33*). Thymosin β4 peptide (Tβ4) (Tb500, PurePeps, tb50010) was dissolved in phosphate-buffered saline solution (PBS, Sigma, P4417-100TAB). A dose of 6mg of the Tβ4 per kg body weight (*34*) was injected intraperitoneally into pregnant mother from E10.5 to E13.5 then embryos were harvested at E14.5. Tamoxifen (Sigma, T5648) was dissolved with 20% ethanol in corn oil (Sigma, C8267) at 20 mg/mL for a stock solution. A dose of 50 mg of the tamoxifen per kg body weight was injected intraperitoneally into the pregnant mother.

### Isolation of cells of the first branchial arch and cells of the trunk

Cells of the first branchial arch (BA1) at E10.5 were harvested according to the protocol we published before (*9, 35*). Briefly, the BA1 or the trunk region at E10.5 was corrected and replaced into ice-cold PBS, then reacted with TrypLE (Gibco, 12605028) for 5 min at 37C. Cells were centrifuged at 100 x g for 5 min at room temperature, then resuspended with PBS. After repeating the centrifuge process again, cells were filtered through a 40 μm cell strainer (Thermo Fischer, 22363547). The isolated single-cell suspension was used for single-cell RNA sequencing or seeded onto Matrigel-coated plate glass for *in situ* hybridization.

### Single-cell RNA sequencing analysis

Single-cell suspensions of the BA1 and the trunk region at E10.5 were harvested as described above. We harvested three of the BA1 at E10.5 from three embryos, placing them into a 1.5 mL microcentrifuge tube to obtain a sufficient number of BA1 cells. The cell number and viability were quantified by using the countess II automated cell counter (ThermoFisher) before loading onto the Chromium Single Cell 3’ v3 microfluidics chip (10× Genomics Inc., Pleasanton, CA). cDNA libraries were sequenced by Illumina Novaseq-6000, generating a total 500 −600 million reads (50,000-70,000 reads per cell). The sequencing data was first pre-processed using the 10× Genomics software Cell Ranger. According to the published protocol, we further developed tSNE-based plots and differential gene expression analysis Filtered feature matrices from Cell Ranger were read into the Seurat R package (*36*). Differential expression in each cluster was determined using the FindMarkers function and the default Wilcoxon test. P-values were corrected using the FDR method. Sexes for each cell in the BA1 of *P0-Cre;caAcvr1* mice at E10.5 was determined based on *Xist* expression and 4 Y-linked gene expressions that had non-zero expression in the cells (*Uty*, *Ddx3y*, *Eif2s3y*, and *Kdm5d*). We used a heuristic such that Y-linked expression < 0.1 and *Xist* expression > 0.8 were considered Female, and the rest were considered Male. To compare high/low *Xist* in Males and Females, we used the MAST test for differential expression (*37*). For all comparisons, a gene is considered differentially expressed if p_val_adj < 0.05 and avg_logFC > 0.1375 (equivalent to a linear fold-change of 1.1).

### Histological analysis

All embryos were dissected and fixed with 4% PFA in PBS at 4 for O/N. Embryos were transferred into 30% sucrose in PBS until embryos were sunk down. Embryos were embedded into OCT-compound (ThermoFisher Scientific, 23-730-571) and 10 or 12 µm of cryo-sections were prepared by a cryostat (Leica CM1950). According to the standard protocol, we stained sections with hematoxylin and eosin, or alcian blue (*9*).

### In situ hybridization and immunofluorescence

*Xist* RNA was detected by using RNA scope Multiplex Fluorescent v2 (ACDBio, 323110) and RNA probe for mouse *Xist* (ACDBio, 454231), according to the standard protocol. Immunohistochemistry was performed as previously described (*9, 38, 39*). Primary antibodies used in this study are the following: SOX9 (1:200 dilution, Millipore, AB5535), GFP (1:200 dilution, R&D, AF4240), Tmsb4x (1:200 dilution, Proteintech, 19850-1-AP), COL2 (1:200 dilution, Abcam, ab34712), and H3K27me3 (Active Motif, 39055). Fluorescence images were taken by confocal microscopy (Nikon, Eclipse Ti).

### RNA preparation and quantitative real-time PCR

Total RNA from the first branchial arch at E10.5 was isolated by using Trizol (Invitrogen, 15596026) followed by ethanol precipitation to purify it. A total of 0.5 μg of total RNA was reverse transcribed using a SuperScript™ II Reverse Transcriptase kit (Invitrogen, 18064022). The amplification was quantified by using Applied Biosystems™ 7500 Real-Time PCR System (Applied Biosystems). Amplified products were monitored using SYBR™ Green PCR Master Mix (Applied Biosystems, 4367659). The following primers were used: mouse *Xist* F 5′-GGTTCTCTCTCCAGAAGCTAGGAAAG −3′; mouse *Xist* R 5′-TGGTAGATGGCATTGTGTATTATATGG −3′; mouse Gapdh F 5′-AGGTCGGTGTGAACGGATTTG-3′; and mouse Gapdh R 5′-AGGTCGGTGTGAACGGATTTG −3′.

### Whole alcian blue staining

Embryos at E14.5 were collected and fixed in Bouin’s solution for 2 hours at room temperature, rinsed with a solution of 1% NH4OH diluted in 70% ethanol until embryos appeared white. Embryos were then equilibrated with 5% acetic acid in 70% ethanol, and stained with 0.05% Alcian blue (Sigma, A5268) in 70% ethanol for 4 hours. After that, embryos were rinsed with 5% acetic acid for 1 hour twice, dehydrated with 100% methanol for 1 hour twice, and cleared in benzyl alcohol benzyl benzoate solution before taking pictures under a stereomicroscope.

### Induction of chondrogenic differentiation for O9-1 cells

All cell cultures were performed in humidified 5% CO2-95% air at 37C. O9-1 cells were obtained from Dr. Robert E. Maxson at the University of Southern California (*3*). We harvested conditioned media from STO cells, which were obtained from Dr. Vesa M. Kaartinen at the University of Michigan, and the conditioned media containing FGF2 (25ng/mL, R&D, 233-FB) and LIF (1,000 U/mL, Millipore, ESG1106) were used for O9-1 cell culture. We induced chondrogenic differentiation of O9-1 cells according to the protocol (*3*). Briefly, O9-1 cells (2.0 x 105 cells/cm2) were seeded into a matrigel-coated plastic culture plate and culture with the conditioned media without FGF2 and LIF for 24 hours. Cells were then cultured with osteogenic differentiation media that are MEM alpha (Gibco, 2185803) containing 10% fetal bovine serum (FBS, Denvile), 1% penicillin/streptomycin (Gibco, 15140-122), Ascorbic acid (50 µg/mL, Gibco, 15140-122), 10 mM β-glycerophosphate, 0.1 µM dexamethazone, and recombinant BMP2 proteins (100ng/mL, R&D, 355-BM) for three days without changing the medium. Cells were trypsinized and harvested to a 15 mL tube, then centrifuged at 100G for 10 min. The supernatant was removed from the tube, and carefully add chondrogenic media that are MEM alpha containing containing 5% fetal bovine serum, 1% Insulin-Transferrin-Selenium (ITS, Gibco, 41400045), 1% penicillin/streptomycin (Gibco, 15140-122), 0.1 µM dexamethasone, sodium pyruvate (1 mM, Gibco, 11360070), recombinant TGF-β3 proteins (10 ng/mL, R&D, 243-B3), and recombinant BMP2 proteins (100ng/mL, R&D, 355-BM). Chondrogenic media were changed every day for 14 days. For the Tβ4 treatment, chondrogenic differentiation media with PBS (as a vehicle condition) or Tβ4 (100 ng/mL, PurePeps, tb50010) were changed every day. After 14 days, pellets were fixed with 4% PFA for 30 min at room temperature, and then prepared cryosections as described above.

### *Ex vivo* culture for the forelimb

The *ex vivo* culture for the forelimb was conducted according to a published protocol (*40*). Briefly, the forelimb of embryos at E10.5 was isolated and placed on a sterile membrane filter (0.4 µm pore size, hydrophilic polycarbonate membrane, Millipore, HTTP02500). The forelimb on the membrane was further placed on a sterile steel mesh and cultured with BGJb media (Gibco, 12591038) containing 1% penicillin/streptomycin (Gibco, 15140-122), and 0.1 µg/mL ascorbic acid (SIGMA-ALDRICH, A4403) with DMSO (as vehicle) or 10 µM X1 compound (GLPBIO, GC26082) in organ culture dish (Falcon, 353037) for 4-5 days. The fluorescence of *HoxA11-GFP* was monitored daily under live imaging with a fluorescent stereomicroscope (Leica M165 FC). The cultured tissues were fixed with 4% PFA for 30 min at room temperature, and then cryosections were prepared by a cryostat ((Leica CM1950).

### Statistical analyses

Statistical analyses were performed by using GraphPad Prism 10.0 software. Data are presented as mean ± standard deviation (SD). Student’s t-test was used for comparison of two groups, and One-way ANOVA with Tukey’s test was used for multiple comparison were performed. All statistical tests were two-sided. P-value is indicated with asterisks; *, p<0.05, and **, p<0.01.

## Acknowledgments

We acknowledge support from the Advanced Genomics Core and the Bioinformatics Core at the University of Michigan Medical School’s Biomedical Research Core Facilities, especially Raymond Cavalcante and Jingqun Ma. We thank Drs. Ken-ichi Yamamura for providing *P0-Cre* mice, Deneen Wellik for HoxA11-GFP mice, and Shigeki Iwase and Yumie Nakamura for providing *Kdm5c^flox/flox^* mice and technical assistance

## Funding

National Institute of Dental and Craniofacial Research (to Y.M.)

1R01DE020843, 1R01DE027662, 1R01HL162939, and 1R21HD119475

## Author contributions

Conceptualization: H.U., Y.M.

Data acquisition: H.U., H.P., S. Khehra., Y.M.

Visualization: H.U., S. Khehra, Y.M.

Data interpretation: H.U., S. Kalantry, Y.M.

Supervision: Y.M.

Writing – original draft: H.U., Y.M.

Writing – review & editing: H.U., H.P., S. Khehra., S. Kalantry, Y.M.

## Data availability

Transcriptomic data from single-cell RNA sequencing have been deposited at the Gene Expression Omnibus (GEO) under accession numbers GSE265974.

