## Supplemental Figures for "X chromosome inactivation as a novel mechanism for chondrogenesis"

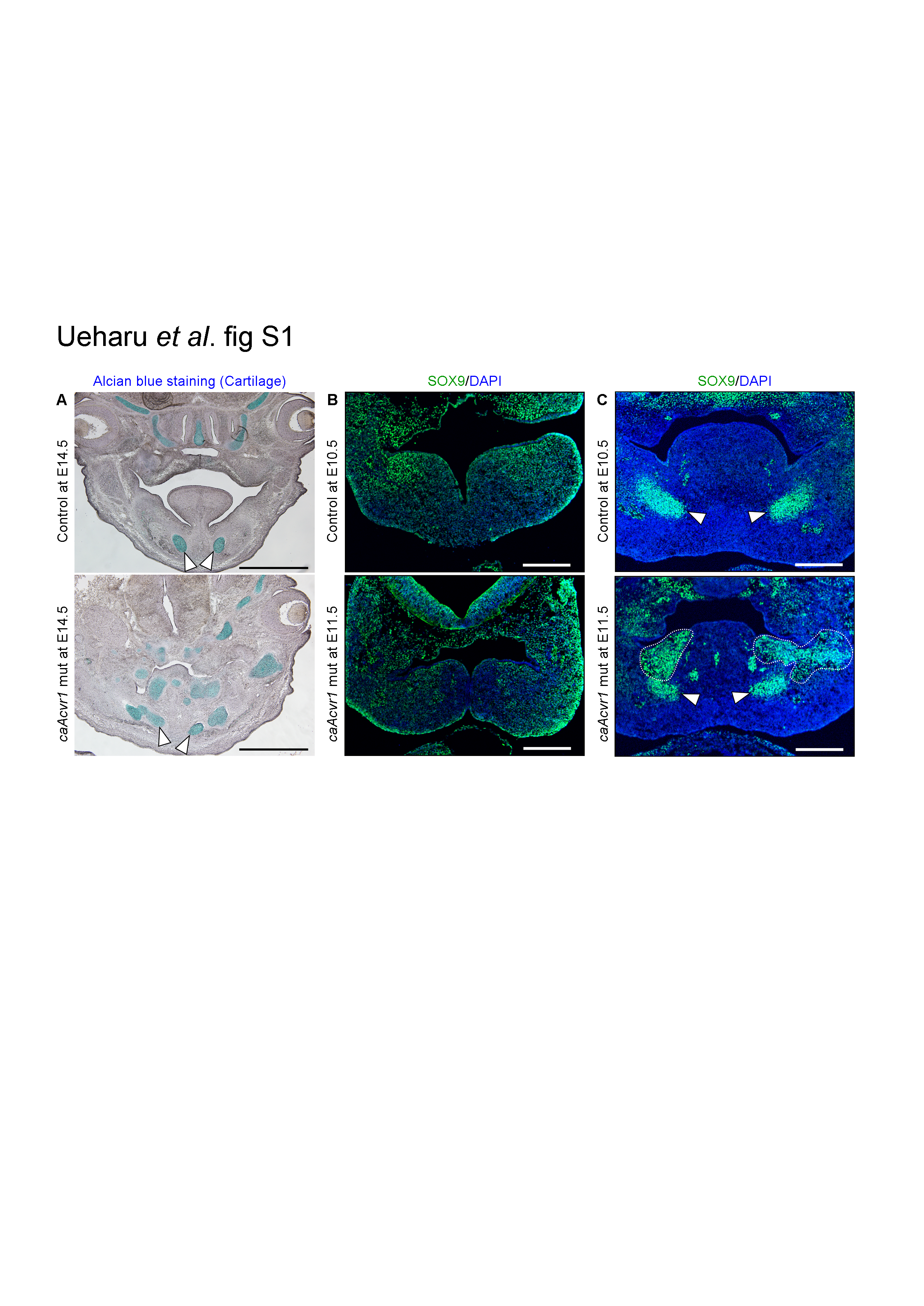
Fig. S1. Evaluation of ectopic cartilage formation during embryonic stage. (A) Representative image of alcian blue staining followed by hematoxylin staining for the frontal sections of *caAcvr1* mice (control) and *P0-Cre;caAcvr1* mice at E14.5 were shown. n=5. (B, C) The localizations of SOX9 in the first branchial arch of control mice and *P0-Cre;caAcvr1* mice at E10.5 (B) and E11.5 (C) were detected by immunohistochemistry for SOX9 (Alexa 488, green). Nuclei were visualized with DAPI (blue). Arrowheads indicated the Meckel’s cartilage. Dotted area indicated ectopic SOX9 aggregation in *P0-Cre;caAcvr1* mice. n=5. Scale; 1 mm (A) and 100 µm (B, C).

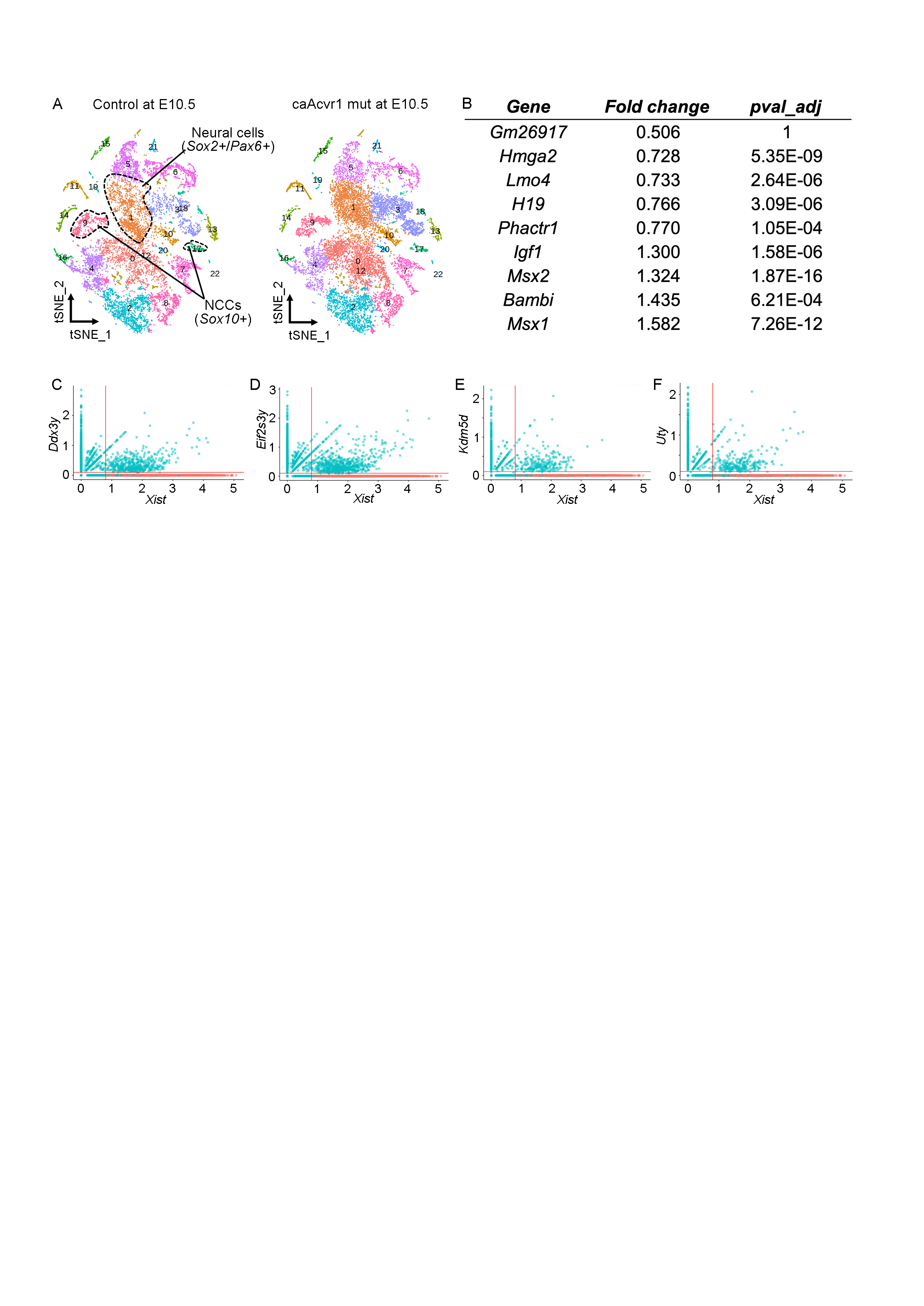

Fig. S2. Single-cell characterization of cells of the trunk region at E10.5 (A-B) and competitive analyses of gene expression in the craniofacial region at E10.5 (C-F). (A) Comprehensive gene expression profiles of cells of the trunk region from *caAcvr1* mice (control) and *P0-Cre;caAcvr1* mice at E10.5 were visualized by t-SNE-based analysis. The dotted area represented a neural cell cluster (Cluster 1; *Sox2*+/*Pax6*+) and neural crest cell clusters (Cluster 9 and 17; *SOX10+*). (B) A comparison of the comprehensive gene expression profiles of neural crest cell clusters revealed genes that showed statistically significant changes in *P0-Cre;caAcvr1* mice compared to control mice. (C-F) Comprehensive gene expression profiles of cells of the BA1 at E10.5, obtained from single-cell RNA sequencing, were further characterized to females and males by expression of *Xist*, Y-linked gene *Ddx3y* (C), *Eif2s3y* (D), *Kdm5d* (E), and/or *Uty* (F). Cells expressed any of Y-linked genes or represented relative *Xist* expression levels less than 0.8 were characterized as male cells (blue dots). Cells without any Y-linked gene expressions and represented relative *Xist* expression higher than 0.8 were characterized as female cells (red dots).

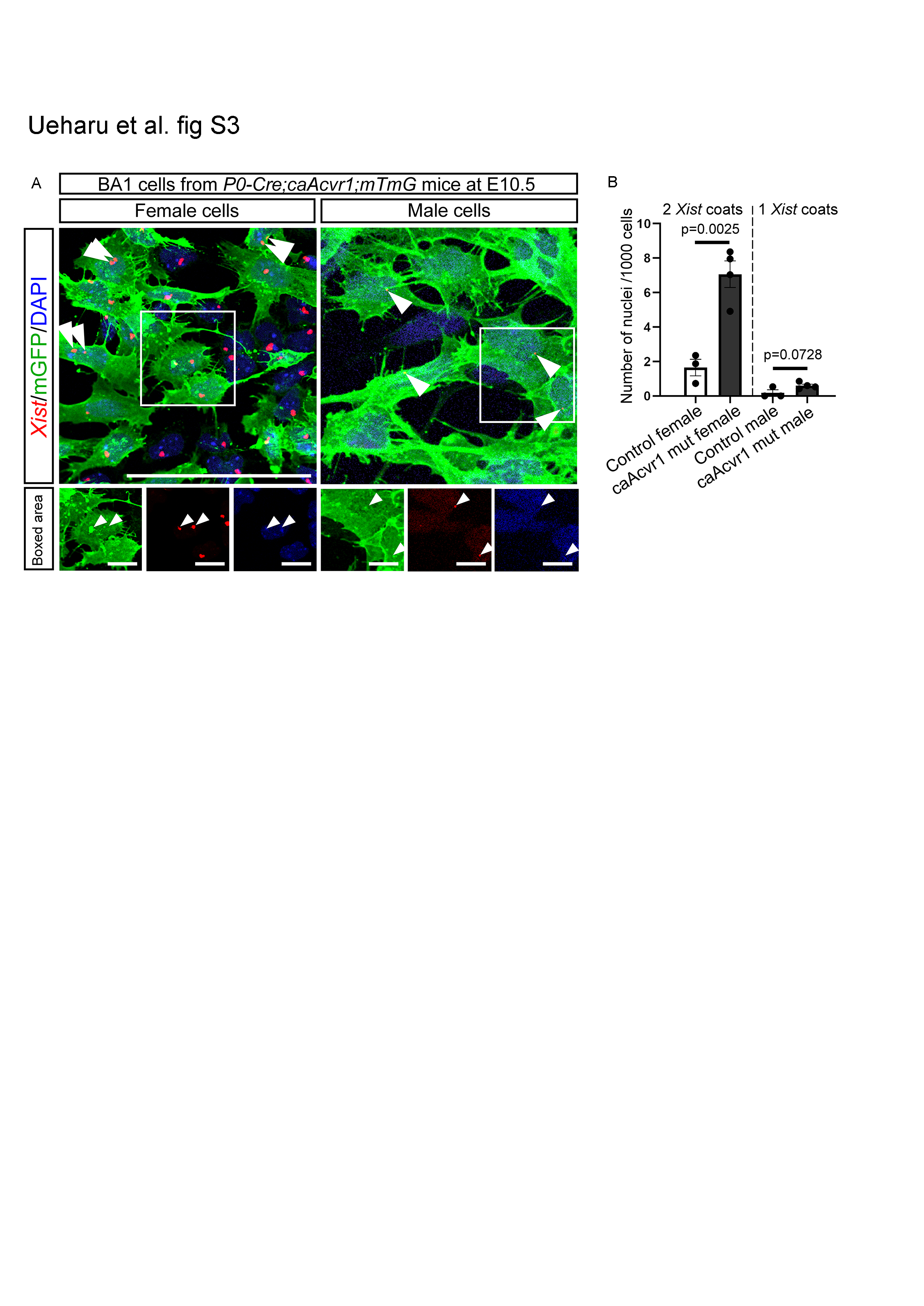

Fig. S3. The presence of ectopic X-inactivation in the cells of the BA1 *in vitro*. (A) Cells of the first branchial arch (BA1) were harvested and cultured onto Matrigel-coated slide glasses for 24 hours. Inactive X-chromosomes were then detected by *in situ* hybridization for *Xist*, followed by immunohistochemistry for GFP to detect plasma cell membrane. The boxed areas were enlarged below, respectively. Arrowheads in the boxed area indicated inactive X-chromosomes. (B) The numbers of cells with two *Xist* RNA coats in females and one *Xist* RNA coat in males were manually counted. Control (n=3) and *P0-Cre;caAcvr1* mice (n=4). Scale; 50 µm (A), 10 µm (A, enlarged). Student’s *t*-test was used for statistical analyses. Each p-value is shown in the figure.

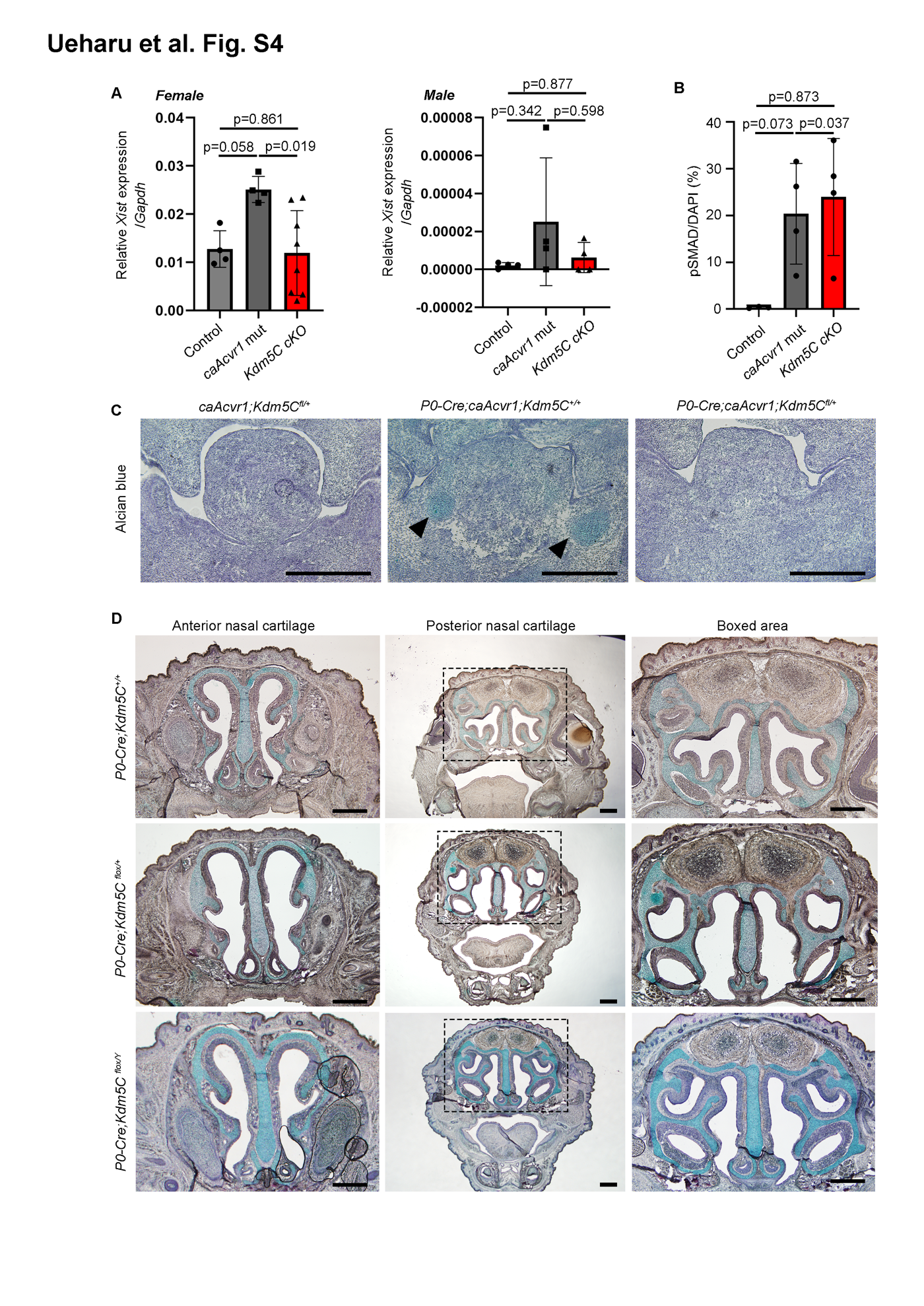

**Fig. S4.** **Analyses for conditional deletion of *Kdm5c* genes in neural crest cells. (A)** Relative *Xist* expression in the first branchial arch (BA1) of *caAcvr1;Kdm5c^fl/+^* female mice (control, n=4), *P0-Cre;caAcvr1;Kdm5c^+/+^* female mice (*caAcvr1* mut, n=4), and *P0-Cre;caAcvr1;Kdm5c^fl/+ or Y^* male mice (*Kdm5c* cKO female, n=8, or *Kdm5c* cKO male, n=4) were shown. The expressions were normalized with the *Gapdh* expression. **(B)** The numbers of cells with phosphoSMAD1/5/9 in *caAcvr1;Kdm5c^fl/+^* mice (control, n=4), *P0-Cre;caAcvr1;Kdm5c^+/+^* mice (*caAcvr1* mut, n=4), and *P0-Cre;caAcvr1;Kdm5c^fl/+^* mice (*Kdm5c* cKO mice, n=4) were shown. **(C)** Representative images of alcian blue staining followed by hematoxylin staining for control mice, *caAcvr1* mut mice, and *Kdm5c* cKO het mice at E13.5 were shown. Arrowheads indicated the ectopic deposition of glycosaminoglycans. n=3 for each genotype. **(D)** Representative images for alcian blue staining followed by hematoxylin staining of *P0-Cre;Kdm5c^+/+^* female mice, *P0-Cre;Kdm5c^fl/+^* female mice, and *P0-Cre;caAcvr1;Kdm5c^fl/+ or Y^* male mice at newborn were shown. n=3 for each genotype. Scale; 100 µm (**C**), 500 µm (**D**).

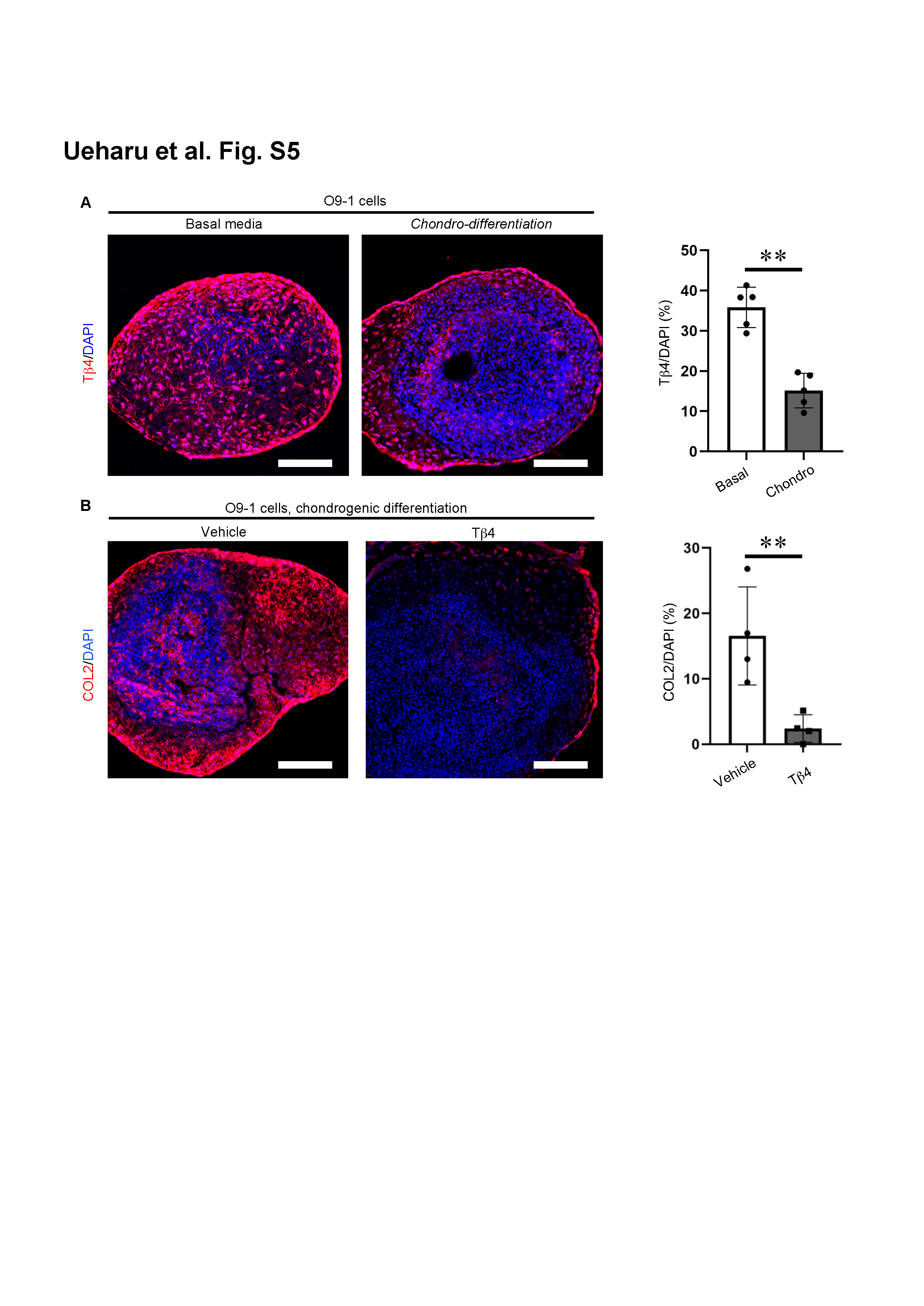

Fig. S5. Involvement of Tβ4 during chondrogenic differentiation *in vitro*. (A) Expressions of Tβ4 in O9-1 cells during chondrogenic differentiation were shown. O9-1 cells were cultured with basal media, which are conditioned media with FGF2 and LIF, or chondrogenic differentiation media. Expression of Tβ4 was detected by immunohistochemistry for Tβ4 (Alexa 594, red). Nuclei were visualized with DAPI (blue). The numbers of Tβ4-expressing cells in each condition were manually counted. n=5. (B) O9-1 cells were cultured with chondrogenic differentiation media including PBS (vehicle) or Tβ4 (100 ng/mL). Chondrogenic differentiation capability was evaluated by immunohistochemistry for COL2 (Alexa 594, red). The numbers of COL2-expressing cells in both conditions were manually counted. n=4. Scale bars; 100 µm. Student’s *t*-test were used for statistical analyses. *=p<0.05, **=p<0.01, N.S.=no significance.

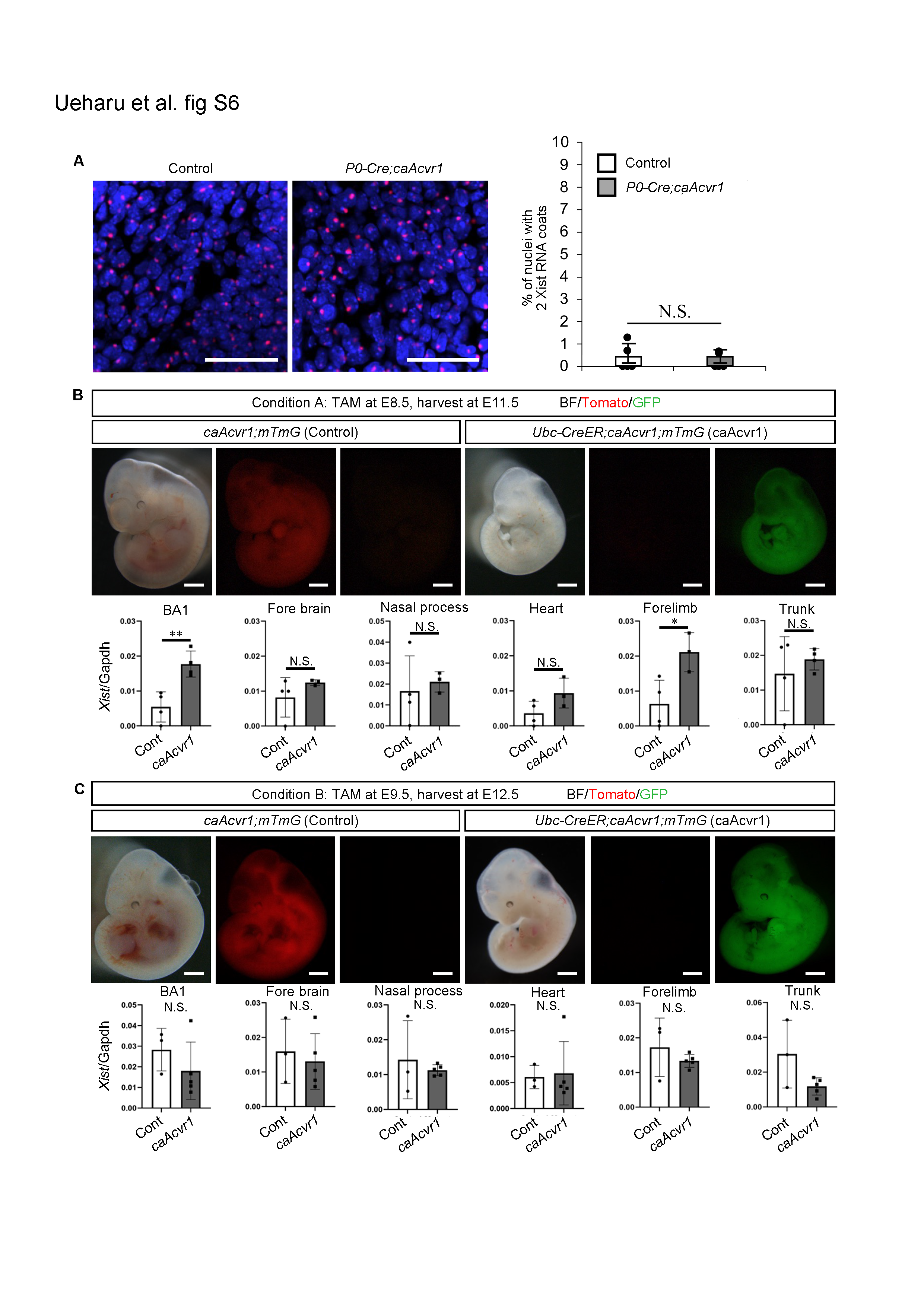

**Fig. S6.** **Spatial and temporal regulation for *Xist* expression by BMP signaling.** **(A)** Inactive X-chromosomes in the dorsal root ganglion at E10.5 were detected by *in situ* hybridization for *Xist* RNA. Nuclei with two *Xist* RNA coats in the trunk region of control female mice (white bar) and *P0-Cre;caAcvr1* female mice (gray bar) were manually counted. **(B)** BMP signaling in *Ubc-CreER;caAcvr1;mTmG* mice was ubiquitously activated by tamoxifen (TAM, 50mg/kg B.W.) from embryonic day 8.5 (E8.5) to E11.5. Then, *Xist* expression in the first branchial arch (BA1), the forebrain, the nasal process, the heart, the forelimb, and the trunk region was quantified by qPCR, respectively. The expression level of *Xist* in *caAcvr1;mTmG* mice (Control) and *Ubc-CreER;caAcvr1;mTmG* mice (*caAcvr1*) were shown. n=3 or 4. **(C)** BMP signaling was ubiquitously activated by tamoxifen from E9.5 to E12.5, then the expression of *Xist* in each tissue was quantified by qPCR. Scale bars; 50 µm (**A**) and 1 mm (**B** and **C**). Student’s *t*-test was used for statistical analyses. *=p<0.05, **=p<0.01, N.S.=no significant difference.

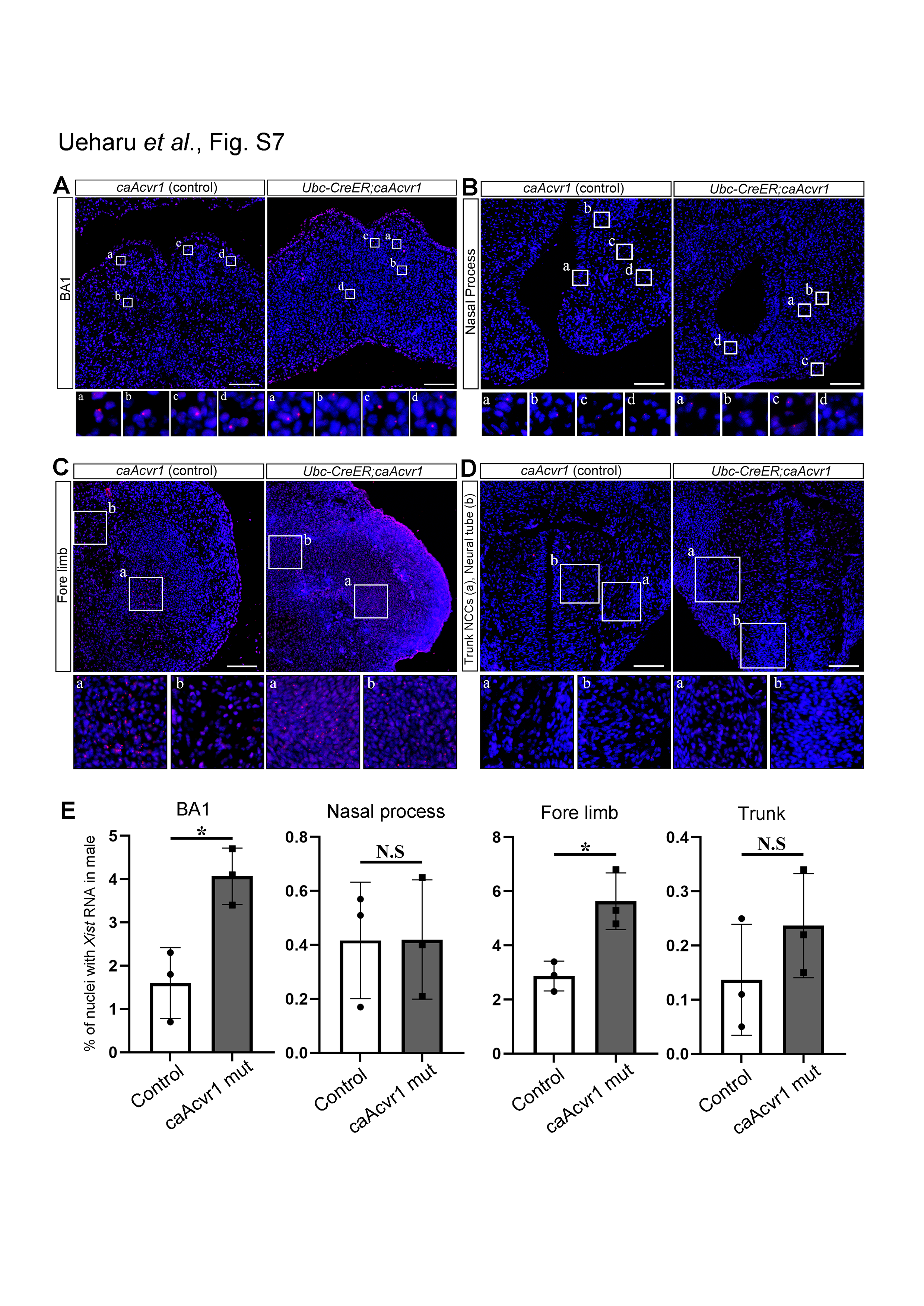
**Fig. S7.** **BMP signaling induced ectopic X-inactivation in a spatial manner. (A-D)** BMP signaling in *Ubc-CreER;caAcvr1* mice was ubiquitously activated by tamoxifen-inducible Cre recombination at E8.5, then embryos were harvested at E11.5. *Xist* RNA in the first branchial arch, the nasal process, the forelimb, and the trunk region of *caAcvr1* male mice (control) and *Ubc-CreER;caAcvr1* male mice was detected by *in situ* hybridization for *Xist* (Opal 570, red), followed by visualization of nuclei by DAPI (blue). **(E)** The numbers of cells with *Xist* RNA in male mice were manually counted. n=3 for each condition. Scale bars; 100 µm. Student’s *t*-test was used for statistical analyses. *=p<0.05, **=p<0.01, N.S.=no significance

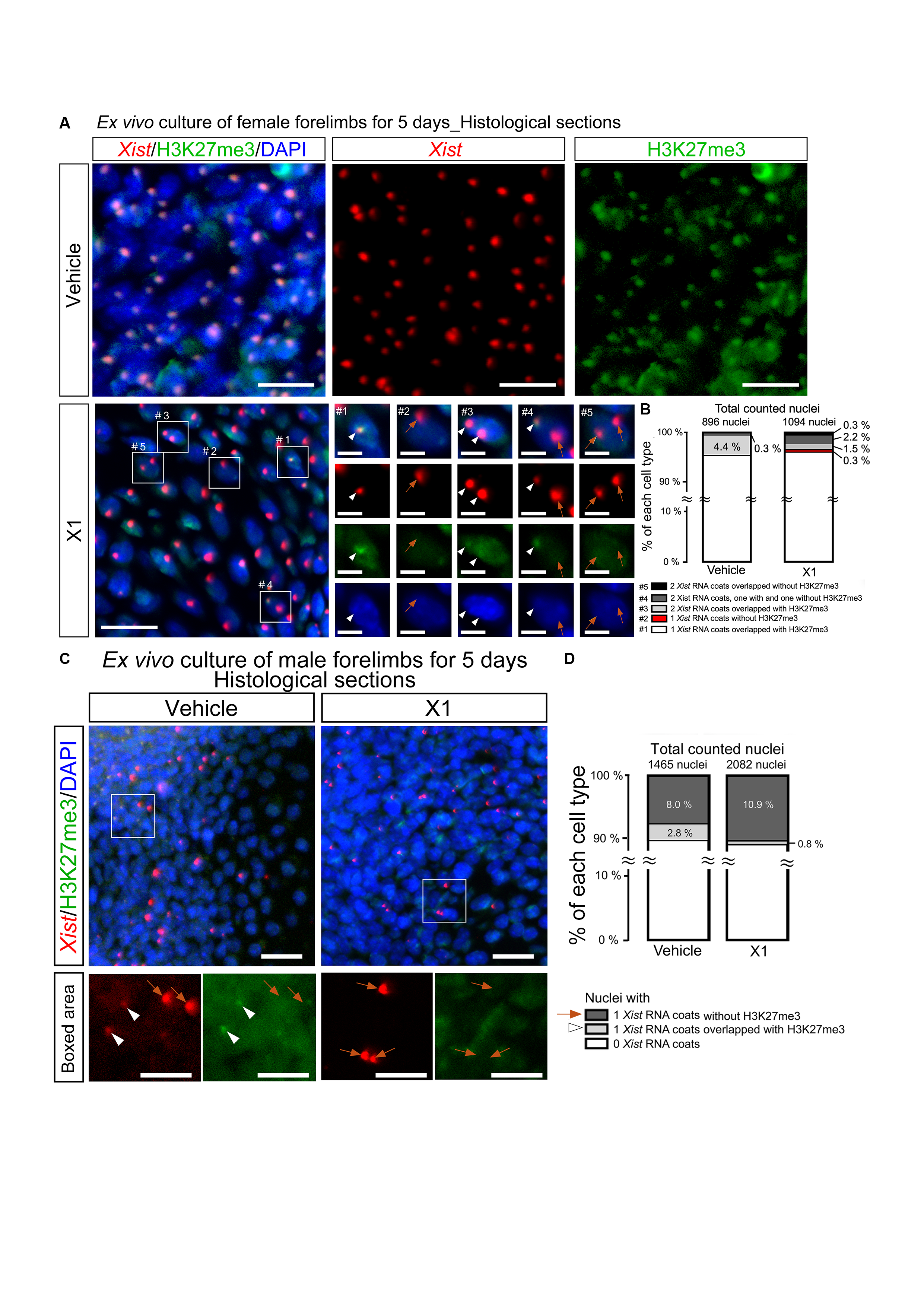
 **Fig. S8.** **Impacts of X1 compound on X-inactivation during forelimb culture.** **(A)** Histological sections of female forelimbs cultured with vehicle or X1 compound for 5 days were used for the detection of *Xist* RNA (red) and H3K27me3 (green) by RNAscope *in situ* hybridization followed by immunohistochemistry. DAPI (blue) was used for nuclei staining. White arrowheads in enlarged pictures indicated *Xist* RNA coats overlapped with H3K27me3, and orange arrowheads indicated *Xist* RNA coats without H3K27me3. **(B)** Percentages of nuclei with: (#1) 1 *Xist* RNA coats overlapped with H3K27me3 (white bar), (#2) 1 *Xist* RNA coats without H3K27me3 (red bar), (#3) *2Xist* RNA coats overlapped with H3K27me3 (light gray bar), (#4) *2Xist* RNA coats, one with and one without H3K27me3 (dark gray bar), and (#5) 2 *Xist* RNA coats without H3K27me3 (black bar), were shown, respectively. A total of 896 nuclei in the vehicle treated forelimbs and a total of 1094 nuclei in the X1 treated forelimbs and were counted. Scale bars; 20 um (low magnification) and 5 um (boxed area). **(C)** Detection of *Xist* RNA (red) and H3K27me3 (green) in male forelimbs cultured for 5 days. DAPI (blue) was used for nuclei staining. White arrowheads indicated nuclei with *Xist* RNA coats overlapped with H3K27me3, and orange arrows indicated nuclei with *Xist* RNA coats overlapped without H3K27me3. Scale bar: 20 um (low magnification) and 10 um (boxed area). **(D)** Percentages of nuclei with 0 *Xist* RNA coats (white), 1 *Xist* RNA coats overlapped with H3K27me3 (light gray), and 1 *Xist* RNA coats without H3K27me3 (dark gray) were shown. A total of 1465 nuclei in vehicle treated forelimbs and a total of 2082 nuclei in X1 treated forelimbs were counted.

**Movie S1**. **A Z-stack image series of a nucleus with 2 *Xist* RNA dots in a cryosection.** The nucleus in the boxed area in **Fig. 2A** was imaged by confocal microscopy using z-stack acquisition, and the consecutive z-stack sections are presented as a movie.

**Movie S2. Three-dimensional reconstruction of the nucleus with 2 Xist RNA dots.** The nucleus in the boxed area in **Fig. 2A** was reconstructed into a three-dimensional image using the z-stack series in **Movie 1**. *Xist* was visualized with red, and membraneGFP was visualized with yellow.

**Movie S3. A Z-stack image of a nucleus with 2 *Xist* RNA dots *in vitro*.** The nucleus in the boxed area in **Fig. S3A** was imaged by confocal microscopy using z-stack acquisition, and the consecutive z-stack sections are presented as a movie.
